# DeSPA uncovers genes with cross-state spatial pattern remodeling and coordinated spatial programs across multi-slice spatial transcriptomics atlases

**DOI:** 10.64898/2026.10.04.756473

**Authors:** Hao Wen, Qunlun Shen, Zhiyuan Yuan, Shuqin Zhang

## Abstract

Spatial transcriptomics increasingly generates multi-slice atlases spanning developmental, regenerative, and disease-associated states. Identifying genes with spatial pattern remodeling across these states is essential for understanding tissue reorganization and the underlying gene programs, yet few existing methods systematically detect such changes while accounting for variation among replicate slices. Here we introduce DeSPA, a statistical framework that models gene specific spatial variation within each slice using a linear mixed model, tests for cross-state remodeling by aggregating evidence across replicate slices, and organizes identified genes into interpretable spatial co-expression modules that reveal coordinated spatial programs. Simulations show that DeSPA achieves high power while controlling the false discovery rate under inter-slice heterogeneity. In mouse embryogenesis, it recovers programs linked to extracellular matrix organization, muscle formation, neural development, and erythrocyte development. In diabetic testis, it reveals disruption of a late-spermatid spatial program consistent with known spermatogenic dysfunction, while maintaining stable gene rankings under coordinate rotation. In axolotl brain, regeneration preserves a developmental like biosynthetic backbone but reorganizes it into regeneration-specific modules. Across two mouse intracerebral hemorrhage models, DeSPA resolves a conserved spatial injury program alongside model-specific synaptic and tissue-response programs. Together, these findings establish DeSPA as a scalable and interpretable framework for detecting spatial pattern remodeling and resolving coordinated spatial programs across diverse biological states.

## 1 Introduction

Spatial transcriptomics links gene expression measurements to their spatial coordinates within tissue sections, allowing molecular variation to be interpreted in its native anatomical context [1, 2]. As spatial platforms have increased in resolution, field of view, and throughput, experimental designs have expanded from individual sections to multi-slice atlases spanning developmental, regenerative and disease-associated states [3–8]. These atlases shift the analytical focus from characterizing spatial organization within single tissue sections to comparative studies of spatial gene expression patterns across different states.

A central task in spatial transcriptomics is the detection of spatially variable genes (SVGs), defined as genes whose expression is structured by spatial location. Early frameworks such as SpatialDE and trendsceek established statistical approaches for identifying such spatial structure [9, 10]. Subsequent methods improved sensitivity, scalability and model flexibility using spatial covariance models, non-parametric statistics, self-organizing maps, diffusion-based representations and scalable Gaussian-process approximations [11–15]. These tools have provided a rich foundation for detecting spatial organization within individual tissue sections.

Multi-slice spatial transcriptomic atlases, however, introduce a distinct comparative problem: identifying genes whose spatial expression patterns are remodeled across multiple biological states. This question is biologically consequential, not merely technically motivated. In mouse organogenesis atlases, spatially resolved expression dynamics have been shown to inform cell-fate specification and tissue-domain formation [5]; injury time-course maps of axolotl telencephalon regeneration have uncovered wound-localized progenitor populations [6]; and tumor spatial architectures, such as tumor-core versus leading-edge transcriptional programs, have been associated with survival and therapy response [16]. Across development, regeneration, disease and treatment, therefore, important signals often manifest as redistribution, expansion, contraction, or relocalization of expression domains, even when marginal expression levels change only modestly.

Despite their recognized biological importance, few existing computational approaches are specifically designed to identify genes with differential spatial expression patterns (DSEPs) across states. Standard differential expression analysis is ill-suited to this task, as it evaluates changes in marginal abundance between states and thus disregards spatial organization. Conversely, conventional SVG detection methods, such as SpatialDE, SPARK/SPARK-X and Moran’s *I* -based screening, are formulated to identify spatial structure within a single tissue section and do not directly assess whether that structure is remodeled across biological states [9, 11, 12]. A few recent studies have begun tackling related comparative problems. SPADE extends SVG detection to between-group comparisons via Gaussian-process regression [17], but its gene-wise fitting is computationally expensive for high-resolution datasets and does not readily extend to multi-state comparisons. River, in contrast, prioritizes genes predictive of slice or condition labels via supervised multi-slice learning using spatially informed neural representations and attribution-based ranking [18]. While this strategy offers flexible modeling of complex spatial patterns, it does not explicitly decompose expression into spatial and non-spatial components, making it unclear whether prioritized genes genuinely reflect DSEPs or merely shifts in marginal expression. Moreover, it relies on external alignment and lacks a calibrated statistical framework to assign confidence to the identified genes. These limitations motivate a framework that directly estimates within-slice spatial structure and tests its remodeling across slices.

Developing such a framework is technically challenging. First, confounding factors such as batch effects, expression shifts, and slice-specific noise can obscure true remodeling. A valid test must therefore distinguish spatial changes from these influences. Second, the geometry and scale of true DSEPs are rarely known in advance. Remodeling may manifest as localized rearrangements or as broad spatial reorganization, making methods that rely on a single neighborhood graph, kernel, or spatial scale inherently sensitive to the chosen parameters. Third, spatial remodeling analysis cannot be reduced to spatial alignment, as imposing alignment across remodeled tissues risks attenuating or distorting the very signals of interest, particularly given that slices often differ in density, orientation, and tissue geometry. Finally, computational efficiency is essential, as growing numbers of spots, slices, and states demand scalable model fitting without compromising statistical calibration.

To address these challenges, we present DeSPA, a model-based statistical framework for identifying genes whose spatial expression patterns vary across multiple states. DeSPA formulates a linear mixed model to decompose gene expression within each slice into non-spatial fixed effects, spatially structured variation and residual variation, thereby isolating the spatial component targeted by DSEP inference. DeSPA then hierarchically integrates evidence across different spatial kernels and slice-pair contrasts with gene-level false discovery rate control. The use of multiple kernels increases sensitivity to spatial remodeling occurring at different scales, whereas slice-pair-based comparison reduces reliance on perfect anatomical alignment. DeSPA further organizes high-confidence genes into cross-slice spatial co-expression modules, linking differential spatial-pattern signals to interpretable biological programs. We evaluated DeSPA using simulations with known ground truth and spatial transcriptomics datasets spanning development, regeneration, disease perturbation and brain injury. These results establish DeSPA as a scalable statistical framework for identifying genes with cross-state spatial pattern remodeling and resolving their coordinated spatial programs across diverse biological states.

## 2 Results

### 2.1 DeSPA overview

DeSPA is a statistical framework for identifying genes with differential spatial patterns across multiple states, such as disease status, treatment, or developmental stage, and resolving the coordinated spatial programs underlying these changes in spatial transcriptomic data. Given gene expression profiles, spatial coordinates, slice identities, state labels, and optional fixed-effect covariates, DeSPA estimates gene-wise spatial components within each slice and tests whether these components vary across states. The framework compares within-slice spatial structures, rather than aligned point-wise expression, and outputs calibrated gene-level evidence of differential spatial patterning together with cross-slice spatial co-expression modules (Fig. 1).

**Figure 1.**
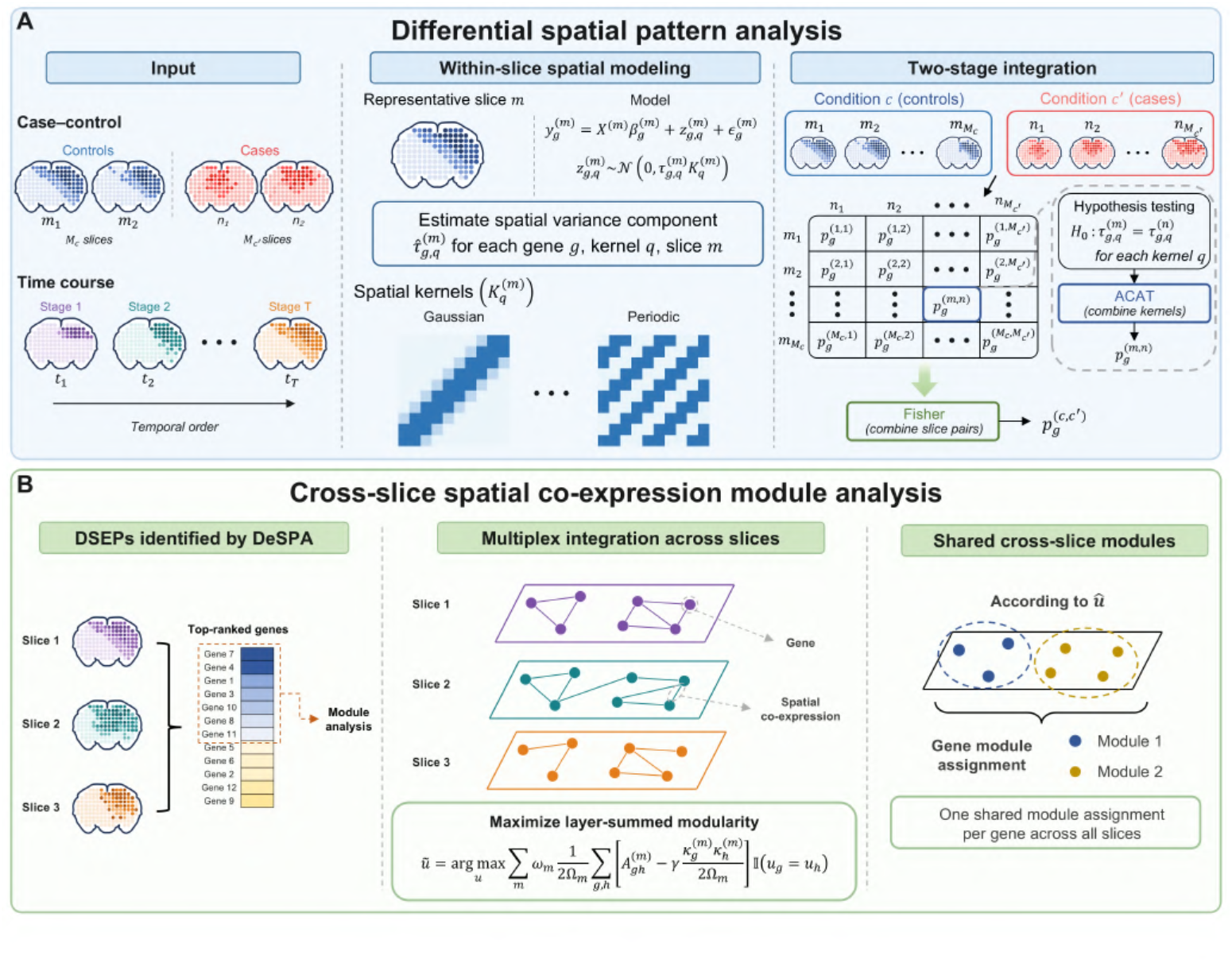
Workflow of DeSPA. **A** Differential spatial pattern analysis. DeSPA accepts case–control or ordered time-course spatial transcriptomic data as input. Within each slice, gene expression is modeled using a linear mixed model that separates fixed-effect covariates, kernel-defined spatial variation and residual noise, and a spatial variance component is estimated for each gene and kernel. For each selected cross-state slice pair, kernel-specific *p*-values are combined using the aggregated Cauchy association test (ACAT). The resulting slice-pair *p*-values are subsequently combined using Fisher’s method to obtain a gene-level *p*-value for the selected comparison set. **B** Cross-slice spatial co-expression module analysis. Top-ranked DSEP genes are used to construct slice-specific spatial co-expression networks, which are treated as layers of a multiplex network. Maximization of the layer-summed modularity objective produces one shared module assignment for each gene across all analyzed slices.

The workflow comprises two connected stages. First, DeSPA performs gene-level testing. For each gene within each slice, a linear mixed model decomposes expression into fixed-effect covariate contributions, spatially structured variation, and residual noise, with spatial structure captured by kernels constructed from within-slice tissue coordinates (Fig. 1A). Since the relevant spatial organization is rarely known in advance, DeSPA evaluates multiple kernels per gene and aggregates evidence across kernels using the aggregated Cauchy association test (ACAT) [19, 20]. Evidence from slices of the same state is then combined via Fisher’s method, and gene-level false discovery rates are controlled using the Benjamini–Hochberg procedure [21]. Computationally, the moment-based parameter estimation reduces the dominant cost to sparse matrix operations, enabling efficient testing across genes, kernels, and slice-pair contrasts. Importantly, because kernels are defined from within-slice distances, DeSPA does not rely on cross-slice coordinate alignment and remains robust to rigid rotations.

In the second stage, DeSPA organizes genes with high-confidence differential spatial patterns into cross-slice spatial co-expression modules that represent coordinated gene programs (Fig. 1B). DeSPA constructs a spatial co-expression network for each slice and integrates these networks into a cross-slice consensus representation to identify modules with coordinated spatial-pattern variation across slices and biological states. These modules provide an interpretable representation of spatial gene programs and differential spatial organization, facilitating downstream biological interpretation through module membership, spatial visualization of member genes and functional enrichment analysis.

DeSPA supports biological analyses of how gene expression programs are spatially reorganized across states. In developmental atlases, it can identify genes whose spatial deployment changes as tissue architecture evolves. In disease or perturbation studies, it can detect genes with divergent spatial patterns, and reveal coordinated gene programs associated with altered tissue organization. In regeneration and injury settings, it can further distinguish recurrent from context-specific spatial programs across slices and biological states. Here, we apply DeSPA to investigate spatial reorganization in four distinct contexts: mouse embryo development, diabetic testis remodeling, axolotl brain regeneration, and mouse intracerebral haemorrhage.

### 2.2 Simulation analysis

We evaluated DeSPA using paired-slice simulations with known ground-truth sets of genes exhibiting differential spatial patterns (**Methods**). Briefly, the simulation design separated slice-level heterogeneity from gene-level spatial-pattern changes. The three slice-level scenarios described how the two slices differed globally: unequal sampling density of spatial locations, a broad slice-specific fixed-effect component in slice 2, or geometric warping of slice 2 coordinates (Fig. 2A). Gene-level settings then defined the status of individual genes (Fig. 2B-D). Null genes had no differential spatial pattern. Interference genes retained the same underlying spatial pattern across slices but were perturbed by either a non-spatial mean shift or residual-noise inflation. Positive genes contained one of five true differential spatial-pattern changes: ring to blob, blob to strip, blob emergence, strip emergence, or increased blob amplitude. Crossing the three slice-level scenarios, two interference settings and five positive settings gave 30 paired-slice benchmarks in total. We compared DeSPA with three competing methods, River, Moran’s *I* screening using both absolute-change and relative-change score (**Methods**), and evaluated power across target FDR thresholds from 0.01 to 0.10.

**Figure 2.**
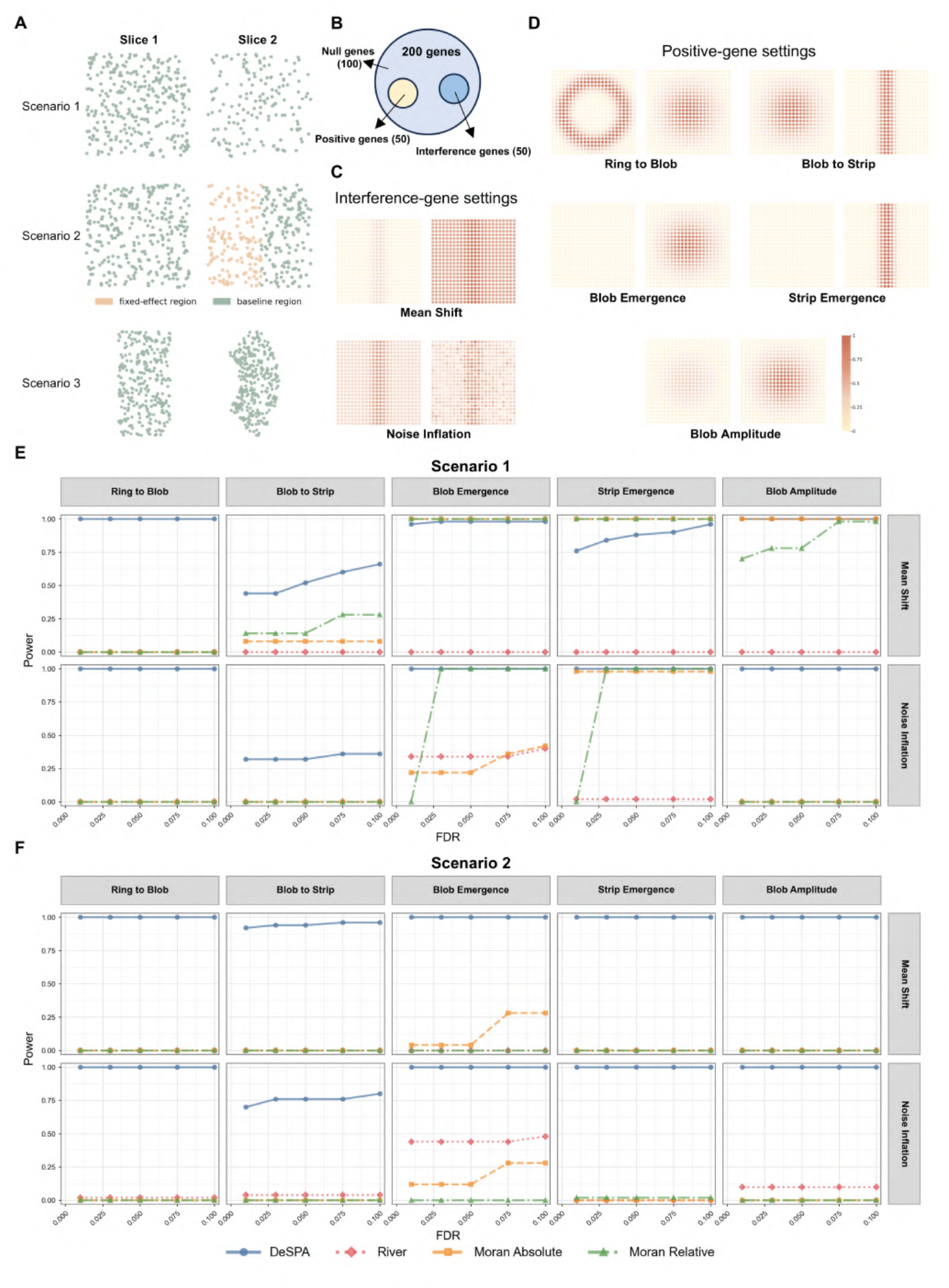
Simulation framework and benchmark performance. **A** Three paired-slice scenarios representing unequal spatial sampling density (Scenario 1), a broad slice-specific fixed-effect region (Scenario 2), and geometric deformation (Scenario 3). **B** Each benchmark contained 200 genes, including 50 positive genes, 50 interference genes and 100 null genes. **C** Interference genes retained the same underlying spatial pattern across slices but were perturbed by either non-spatial mean shifts or residual-noise inflation. **D** Positive genes represented five differential spatial-pattern settings: ring-to-blob, blob-to-strip, blob emergence, strip emergence and blob-amplitude change. **E**,**F** Power curves for Scenario 1 and 2, respectively. Columns correspond to the five positive spatial-pattern settings, and rows correspond to the two interference settings, mean shift and noise inflation. DeSPA showed consistently high power across most simulation settings, including heterogeneous sampling, fixed-effect variation and geometric deformation.

Across all 30 benchmarks, DeSPA showed the most stable overall performance. Its mean power across target FDR thresholds remained between 0.89 and 0.92. At a target FDR of 0.05, DeSPA achieved a mean power of 0.90 across all settings, compared with 0.06 for River, 0.38 and 0.36 for Moran’s *I* screening with both absolute-change and relative-change score, respectively. DeSPA was the top-performing or tied top-performing method in 26 of the 30 benchmarks and achieved power of at least 0.8 in 25 benchmarks. The following detailed analysis is on the results obtained at a target FDR of 0.05 unless otherwise stated.

In scenario 1, where the two slices represented the same tissue region but were sampled at different spatial densities, DeSPA recovered most positive patterns with high power (Fig. 2E). It detected ring-to-blob and blob-amplitude changes with power 1.00 under both interference settings, and detected blob or strip emergence with power 0.88–1.00. The main exception was the blob-to-strip transition, for which power decreased to 0.52 under mean-shift interference and 0.32 under noise-inflation interference. This transition is more challenging because both slices contain spatially structured signals, so the differential signal reflects redistribution of an existing pattern, while unequal sampling density further obscures the resulting boundary. Moran’s *I* screening mainly recovered emergence or amplitude changes, but had little power for topology changes, consistent with global spatial-autocorrelation summaries being more sensitive to the structure strength than to spatial shape. River yielded fewer detections in this density-mismatch scenario. This is likely because River relies on reliable coordinate correspondence between slices, while the relatively sparse spatial locations and unequal sampling densities in our synthetic benchmark may have increased coordinate-matching uncertainty, thereby reducing its apparent detection power.

In scenario 2, slice 2 contained a broad fixed-effect contribution, providing a direct test of whether methods could distinguish true gene-specific spatial-pattern changes from slice-level nuisance variation (Fig. 2F). DeSPA remained highly powered across positive settings, with power ranging from 0.94 to 1.00 under mean-shift interference and from 0.76 to 1.00 under noise-inflation interference. In contrast, Moran’s *I* screening showed limited recovery, with maximum power of 0.12 and 0.02 using absolute-change and relative-change score, respectively. River also produced few detections, except for the blob-emergence setting under noise-inflation interference. These results reflect the distinct statistical targets of the methods: those based on global spatial-variability summaries can be influenced by spatially structured nuisance variation in the negative-control genes, whereas DeSPA retained power by focusing on differential spatial-pattern structure while accounting for fixed-effect contributions.

In scenario 3, spatial patterns were generated in a shared reference coordinate system, with slice 2 subjected to geometric warping (Supplementary Fig. 1). DeSPA again showed high power for most settings, reaching power 1.00 for blob-to-strip, blob emergence and blob-amplitude changes under both interference settings, and 0.96–1.00 for strip emergence. The limitation was the ring-to-blob transition, where power was low under mean-shift interference and moderate under noise-inflation interference. The combination of topology change and coordinate deformation made the annular-to-central redistribution harder to distinguish from non-target variation. Moran’s *I* screening performed well in several mean-shift settings, likely because global auto-correlation is less dependent on pointwise alignment, but their recovery decreased under noise inflation for topology and intensity changes. River again produced fewer detections across most deformed settings, with its strongest recovery observed for the none-to-blob appearance change under residual-noise-inflation interference.

Overall, these simulations show that DeSPA is effective in distinguishing true differential spatial patterns from sampling-density differences, broad slice-specific fixed effects and non-target expression perturbations. Performance was somewhat lower in more challenging settings involving topology changes between spatially variable structures, particularly blob-to-strip transitions under density mismatch, and severe geometric deformation such as the ring-to-blob transition under strong mean-shift interference. These results help define the current operating range of DeSPA and suggest opportunities for future extensions incorporating deformation-aware registration or multi-scale topological features.

### 2.3 DeSPA resolves stage-dependent spatial remodeling programs during mouse embryogenesis

During embryogenesis, spatially restricted gene expression programs are continually reshaped as tissues grow, fold, differentiate and acquire organ-specific identities. Identifying genes whose expression patterns are remodeled across developmental process can therefore reveal molecular programs underlying morphogenesis, lineage specification and tissue regionalization. We applied DeSPA to a Stereo-seq mouse embryo developmental series [5], consisting of eight ordered stages from E9.5 to E16.5, with well-resolved tissue structures across development and 5,913–121,767 spatial locations per stage (Fig. 3A). As embryos change substantially in size and shape across stages, DeSPA avoids direct coordinate registration by comparing stage-specific spatial components, enabling identification of genes with developmental spatial remodeling without one-to-one spatial correspondence.

**Figure 3.**
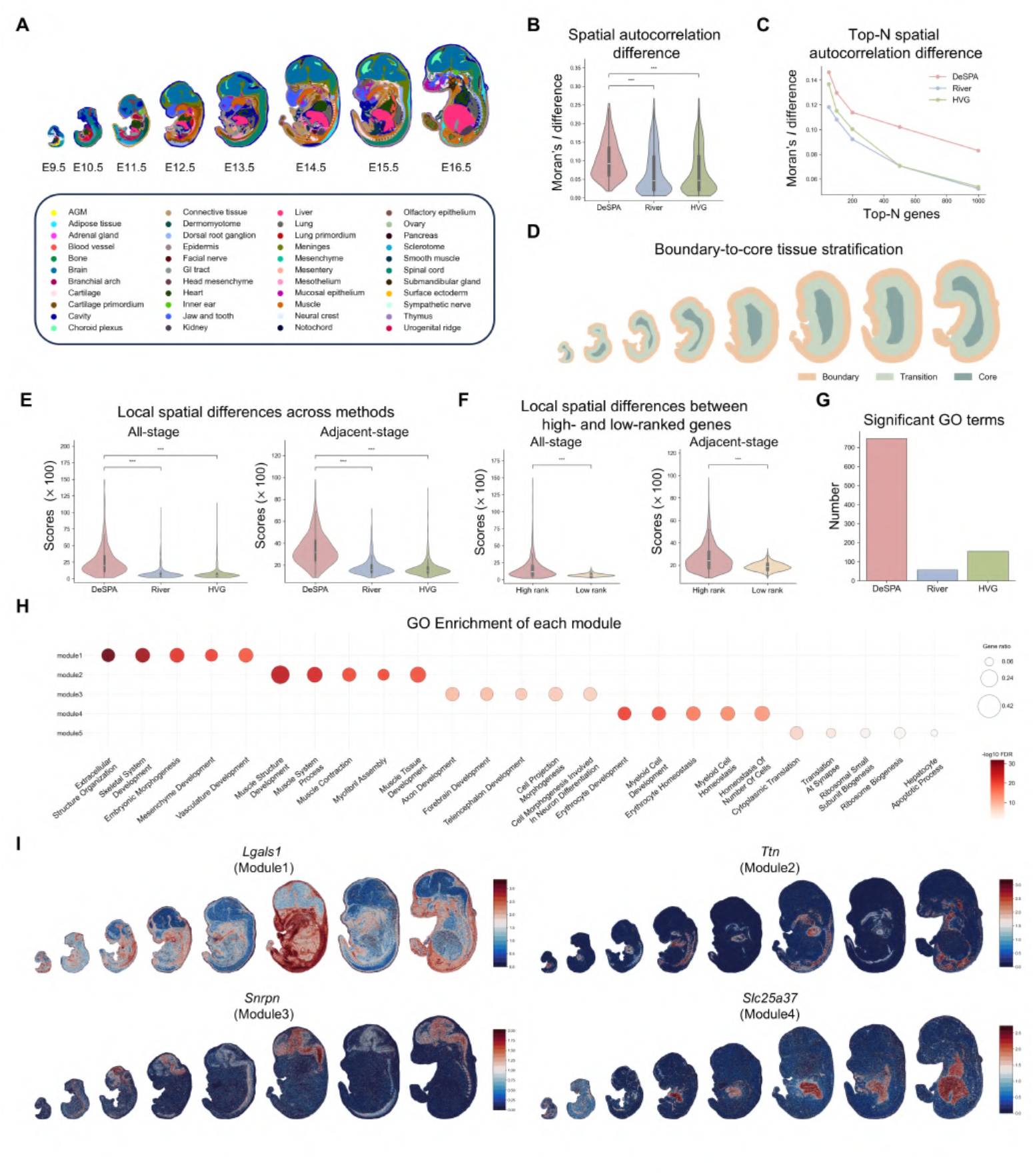
DeSPA resolves stage-dependent spatial remodeling programs during mouse embryogenesis. **A** Spatial annotations of mouse embryo sections from E9.5 to E16.5. **B** Distributions of the adjacent-stage spatial autocorrelation difference scores among the top 500 genes ranked by DeSPA, River and the highly variable gene (HVG) ranking. **C** Mean spatial autocorrelation difference score among the top *N* genes ranked by each method, with *N* = 50, 100, 200, 500 and 1,000. **D** Annotation-independent boundary-to-core tissue stratification used to quantify regional expression redistribution across developmental stages. **E** Distributions of the all-stage and adjacent-stage local spatial pattern difference scores among the top 500 genes ranked by DeSPA, River and HVG. Scores are displayed after multiplication by 100. **F** Distributions of the two local spatial pattern difference scores among the top- and bottom-2,000 DeSPA-ranked genes. **G** Numbers of significantly enriched Gene Ontology (GO) Biological Process terms among the top 500 genes selected by each method. **H** GO Biological Process enrichment of DeSPA-derived cross-slice spatial co-expression modules. **I** Spatial expression patterns of representative genes in each module, including *Lgals1, Ttn, Snrpn* and *Slc25a37*, across embryonic stages.

Genes prioritized by DeSPA showed stronger developmental differences in global spatial autocorrelation. Using the spatial autocorrelation difference score, defined as the mean absolute difference in Moran’s *I* between adjacent developmental stages (**Methods**), the top 500 genes identified by DeSPA achieved a median score of 0.092, approximately twofold higher than River (0.046, *p*-value = 2.30 × 10 ^-26^, two sided Wilcoxon rank-sum tests) and highly variable genes (HVGs; 0.047, *p*-value = 6.04 × 10^-27^) (Fig. 3B). Across the top 50, 100, 200, 500 and 1,000 genes, DeSPA achieved mean spatial autocorrelation difference scores of 0.146, 0.130, 0.114, 0.102 and 0.083, respectively, consistently exceeding those of the compared methods (Fig. 3C). These results demonstrate that DeSPA preferentially ranks genes whose spatial auto-correlation changes across embryonic stages, rather than genes that are merely highly expressed, highly variable or spatially patterned at one stage.

Genes involved in embryonic morphogenesis are expected not only to change their global spatial autocorrelation, but also to shift expression region across local tissue compartments as organs emerge, expand and become spatially refined. We therefore used an annotation-independent boundary-to-core tissue stratification and quantified local spatial pattern differences using all-stage and adjacent-stage local spatial pattern difference scores (Fig. 3D; **Methods**). Higher values indicate stronger redistribution of a gene’s expression across boundary, transition and core regions during development. The top-ranked genes identified by DeSPA showed substantially stronger local spatial pattern differences than genes selected by River or HVG ranking. Among the top 500 genes, DeSPA achieved a mean all-stage local spatial pattern difference score of 31.78, compared with 16.03 for River and 15.21 for HVGs (Fig. 3E). DeSPA also produced the strongest adjacent-stage local spatial pattern difference score, with a mean value of 4.54, compared with 2.29 for River and 2.17 for HVGs. Within the DeSPA ranking, the top 2,000 genes also showed greater local spatial pattern differences than the bottom 2,000 genes (Fig. 3F). These results indicate that DeSPA-ranked genes preferentially undergo dynamic redistribution between boundary, transition and core tissue regions, consistent with spatial programs that are reorganized as embryonic structures form.

These quantitatively defined spatial-remodeling genes corresponded to coherent developmental biology. Gene Ontology enrichment analysis of the top 500 genes revealed a marked functional distinction between DeSPA and the compared methods. The top-ranked genes by DeSPA were enriched for 746 significant biological-process terms, led by muscle structure development (FDR = 9.14 × 10^-29^). By contrast, top ranked genes by River and HVGs were dominated by cytoplasmic translation (FDR = 8.48 × 10^-53^ and 6.54 × 10^-71^, respectively), with 56 and 155 significant terms, respectively (Fig. 3G, Supplementary Fig. 2). This functional shift suggests that DeSPA preferentially recovers genes associated with tissue construction, morphogenesis and developmental remodeling, whereas the compared methods are more strongly influenced by broadly active translational programs.

To investigate whether the genes prioritized by DeSPA could be grouped into biologically interpretable developmental programs, we performed cross-slice spatial co-expression module analysis (Fig. 3H; **Methods**). Module 1 was enriched for extracellular structure organization and contained extracellular-matrix and tissue patterning genes, including *Col5a2* and *Lgals1*, with *Msx1* providing an additional developmental patterning signal [22]. This module is consistent with matrix-associated tissue remodeling during embryogenesis. Module 2 was enriched for muscle structure development and contained canonical muscle-associated genes, including the sarcomeric gene *Ttn* and the skeletal-muscle regulatory light-chain gene *Mylpf*, together with *Nrk* [23–25]. This module therefore marked a spatial program associated with embryonic muscle formation. Module 3 was enriched for neural developmental processes, including axon and forebrain development, and included neural-associated genes such as *Mllt11* and *Fez1* [26, 27]. Together with additional genes including *Snrpn* in this module, these patterns support a spatial program linked to neural differentiation and tissue organization. Module 4 was enriched for erythrocyte development and contained haemoglobin genes, including *Hbb-bs* and *Hbb-bt*, together with the mitochondrial iron transporter *Slc25a37* [28, 29]. This module therefore reflected an erythroid and haematopoietic spatial program during embryogenesis. These results show that DeSPA identifies genes with differential spatial patterns and organizes them into cross-stage modules representing coordinated spatial programs associated with extracellular matrix organization, muscle formation, neural development and erythroid differentiation.

Spatial expression maps of representative genes further supported these module-level interpretations. *Lgals1* showed weak and scattered expression at early stages but became broadly and strongly expressed across large embryonic regions at later stages, consistent with extracellular-matrix-associated tissue remodeling. *Ttn* showed weak spatial expression at early stages and became concentrated in discrete late stage domains, matching the muscle-formation module. *Snrpn* showed stage-dependent enrichment in restricted neural-associated regions, whereas *Slc25a37* became prominent in compact internal domains at later stages, consistent with erythroid and haematopoietic activity (Fig. 3I, Supplementary Fig. 3).

Together, the analysis on the mouse embryogenesis shows that DeSPA prioritizes genes with stronger stage-associated spatial autocorrelation differences, greater local spatial pattern differences and clearer developmental module structure than genes ranked by the comparison methods, supporting its ability to recover biologically interpretable spatial programs during embryogenesis.

### 2.4 DeSPA identifies rotation-invariant disease-associated spatial programs in matched testis sections

Disease-associated tissue dysfunction extends beyond changes in gene expression levels to the spatial organization of expression patterns. This is particularly relevant in the testis, where spermatogenesis depends on the ordered arrangement of germ-cell states within seminiferous tubules. We therefore applied DeSPA to a Slide-seq dataset containing three leptin-deficient diabetic mouse testis sections and three wild-type (WT) sections, analyzed as three fixed one-to-one case–control pairs [7]. Individual sections contained approximately 27,194–42,776 spatial locations and 23,514–24,450 detected genes, and cell-population annotations revealed structured spatial organization across the testis sections (Fig. 4A).

**Figure 4.**
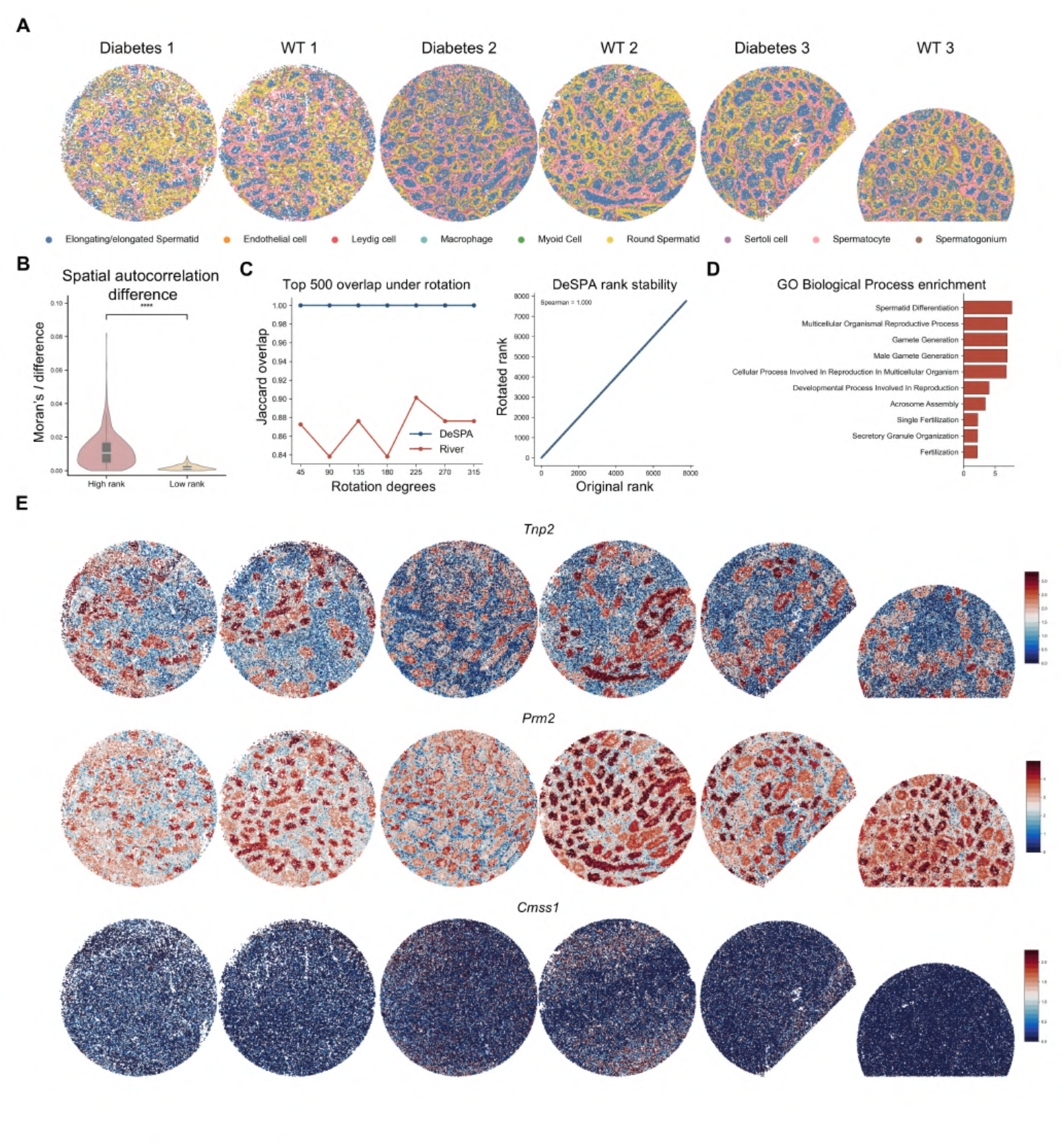
DeSPA identifies robust disease-associated spatial patterns in matched diabetic and wild-type testis sections. **A** Spatial annotations of three diabetic and three matched wild-type (WT) testis sections. **B** Distributions of the absolute diabetic-WT spatial autocorrelation difference scores among the top- and bottom-500 DeSPA-ranked genes. **C** Rotation analysis showing top-500 gene overlap and rank stability after rotating diabetic slices. DeSPA preserved identical rankings under rigid coordinate rotation. **D** GO Biological Process enrichment of DeSPA-ranked genes, highlighting reproductive and spermatogenesis-associated processes. **E** Spatial expression patterns of representative genes, including *Tnp2, Prm2* and *Cmss1*, across diabetic and WT sections.

To assess whether DeSPA ranking reflects state-associated spatial reorganization, we compared the absolute case-control difference in Moran’s *I* between top- and bottom-ranked genes. The top 500 genes exhibited a median absolute difference of 0.0105 between diabetic and WT testes, approximately sevenfold higher than the bottom 500 genes (0.00148) (*p*-value = 6.91 × 10^-107^, two-sided Wilcoxon ranksum test; Fig. 4B). This shows that DeSPA preferentially ranks genes whose spatial autocorrelation differs between diabetic and WT testes.

As these slices exhibit high morphological similarity and are difficult to align, a rotation-invariant method becomes necessary. To evaluate this property, we therefore rigidly rotated the spatial coordinates of each diabetic section by 45**°** increments (45**°** to 315**°**), while preserving the correspondence between each spatial location and its expression profile. Across all rotation angles, the Spearman correlation between the original and rotated DeSPA rankings was 1.00, and the top-500 Jaccard index was also 1.00. By contrast, River achieved Spearman correlations of 0.933–0.977 and top-500 Jaccard indices of 0.838–0.901 under the same rotations. Rank–rank plots showed that DeSPA rankings remained on the diagonal, whereas River rankings displayed visible displacement (Fig. 4C, Supplementary Fig. 4A). These results show that DeSPA was robust to tissue orientation. Importantly, this invariance was accompanied by appropriate control of false discoveries. In same-condition negative-control comparisons across all rotation angles, DeSPA detected no significant genes (Supplementary Fig. 4B). Thus, DeSPA remained both geometrically stable and statistically conservative under rotation, indicating that rigid coordinate transformations alone do not generate spurious differential spatial signals.

Gene Ontology enrichment analysis of the top 500 DeSPA-ranked genes identified 14 significant biological-process terms at FDR *<* 0.05. The most significant term was spermatid differentiation, comprising 49 genes (FDR = 2.06 × 10^-8^; Fig. 4D). Other enriched terms included gamete generation, multicellular organismal reproductive process, acrosome assembly, single fertilization and fertilization. The contributing genes converged on late spermatid biology: *Prm1, Prm2, Tnp1* and *Tnp2* mediate chromatin remodeling and packaging during spermiogenesis [30, 31], whereas *Akap4, Cabyr* and *Fsip2* contribute to flagellar assembly or sperm structural maturation [32, 33]. DeSPA thus recovered a coherent program centered on late spermatid differentiation and sperm maturation, a finding strongly supported by external evidence. In the original analysis of the same Slide-seq dataset, genes predominantly expressed in elongating/elongated spermatids, including *Prm1, Prm2, Odf1* and *Smcp*, were significantly downregulated in *ob/ob* testes, with *Smcp, Odf1* and *Malat1* showing altered spatial expression patterns [7]. Notably, these three spatially altered genes ranked 1st, 14th and 87th, respectively, in the DeSPA analysis. The same study also reported reduced spatial purity of elongating/elongated spermatids in *ob/ob* seminiferous tubules, indicating disrupted spatial organization [7]. Independent studies have further shown fewer spermatogonia, spermatocytes and spermatids, together with an absence of sperm within seminiferous tubules, in leptin-deficient mice [34], as well as altered expression of *Prm1, Prm2, Tnp1* and *Tnp2* in a streptozotocin-induced diabetic model [35]. These observations link the late-spermatid program recovered by DeSPA to well-established features of diabetic spermatogenic dysfunction.

To determine whether the DeSPA prioritized genes constituted one or more spatial programs, we grouped the top-ranked genes into cross-slice spatial co-expression modules. This analysis resolved five modules containing 129, 118, 94, 86 and 73 genes, respectively. Module 3 showed the strongest reproductive annotation, with significant enrichment for cellular process involved in reproduction in multicellular organism (FDR = 2.07×10^-4^), and spermatid differentiation (FDR = 2.03×10^-3^). This module contained sperm structural genes such as *Cabyr, Catsper1, Fsip1* and *Fsip2* (Supplementary Fig. 5), supporting a spatial program associated with sperm maturation rather than a single-gene effect [32, 33, 36]. Module 4 was enriched for response to xenobiotic stimulus and contained stress- or detoxification-associated genes, including *Lcn2, Aldh1a1, Gstm1* and *Gstm2* [37, 38], suggesting a spatial program distinct from the spermatid-maturation axis. Ligand–receptor analysis of the same WT and *ob/ob* spatial resource further revealed spatial redistribution of inflammation-related signals, consistent with stress-associated remodeling of the diabetic testicular microenvironment [39].

**Figure 5.**
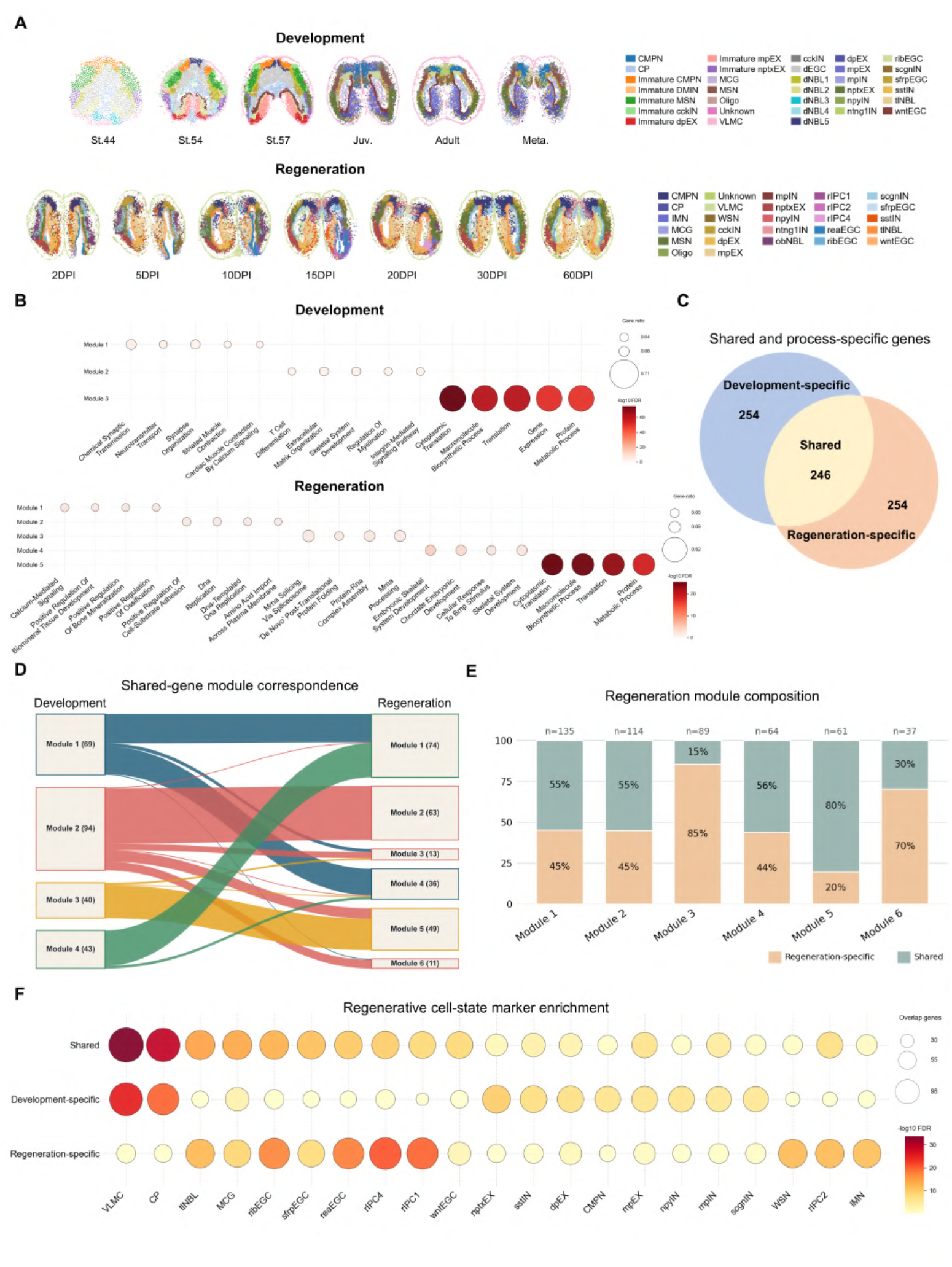
DeSPA distinguishes shared and process-specific spatial programs in axolotl brain development and regeneration. **A** Spatial annotations of axolotl brain sections across development and regeneration. DPI: days post-injury. Cell-type and cell-state abbreviations are defined in Supplementary Table. 2. **B** GO Biological Process enrichment of DeSPA-derived spatial co-expression modules in the developmental and regeneration trajectories. **C** Overlap between the top 500 DeSPA-ranked genes in development and regeneration, separating shared, development-specific and regeneration-specific genes. **D** Alluvial plot showing module correspondence for shared genes between development and regeneration. **E** Composition of regeneration modules according to shared and regeneration-specific gene membership. **F** Enrichment of shared, development-specific and regeneration-specific genes in regenerative cell-state marker sets.

Spatial expression maps of representative genes supported this module-level interpretation. Genes from modules 1 and 2, including *Smcp, Prm2, Odf1, Prm1* and *Tnp2*, displayed spatially organized expression patterns consistent with late spermatid maturation, chromatin condensation and sperm structural organization, even though some corresponding modules showed weak GO enrichment as a whole (Fig. 4E, Supplementary Fig. 6). In contrast, *Lcn2* displayed a distinct spatial pattern consistent with the stress-associated module rather than the sperm-maturation axis (Supplementary Fig. 6). These tissue-level maps indicate that the diabetic-testis signal, rather than being a diffuse collection of disease-associated genes, can be resolved into spatially distinct spermatid-associated and stress-associated programs. Together, these results show that differential spatial-pattern analysis reveals a tissue-level manifestation of diabetic spermatogenic dysfunction: disruption of late-spermatid organization within seminiferous tubules together with a distinct stress-associated program.

**Figure 6.**
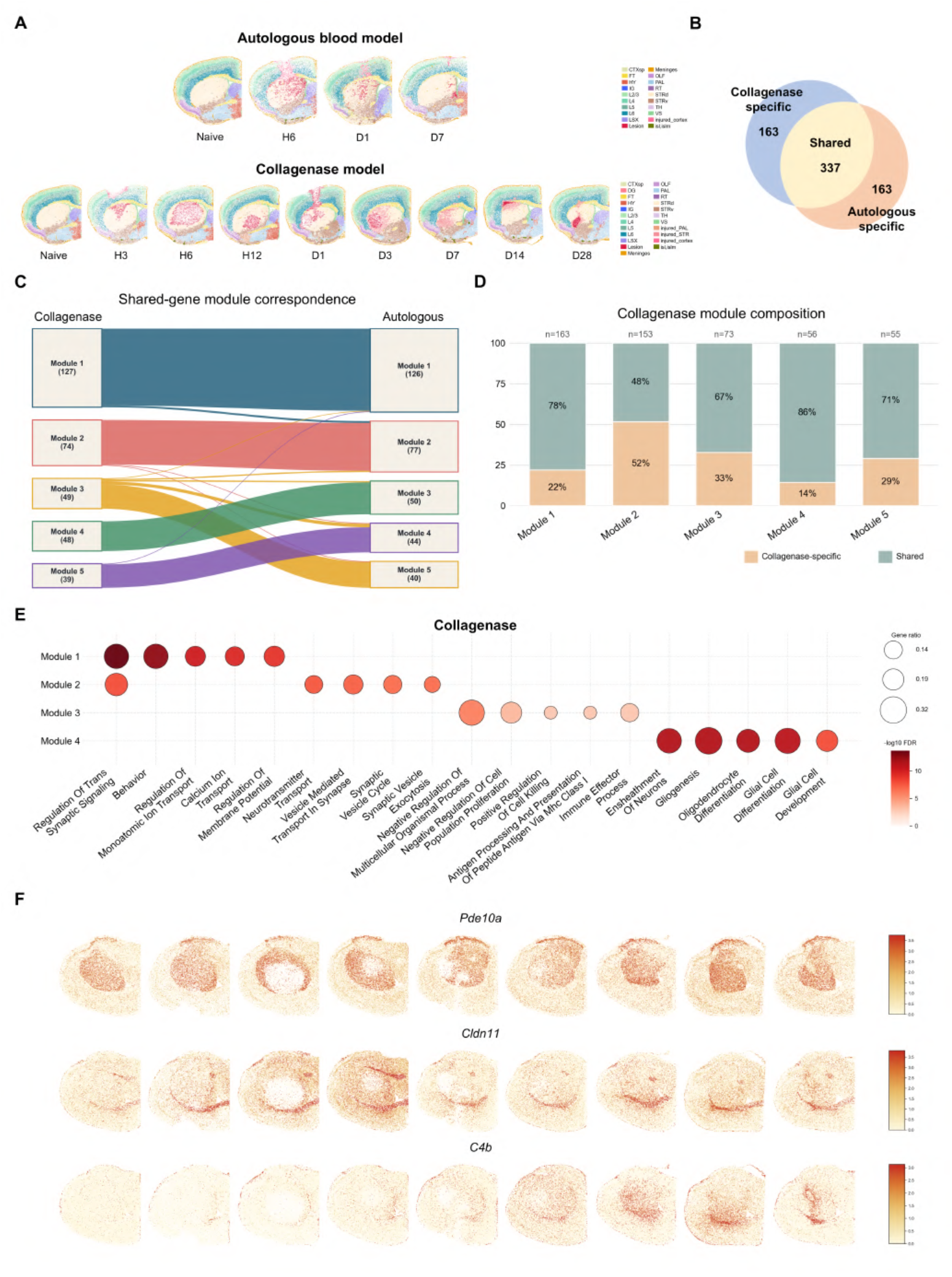
DeSPA resolves conserved and model-biased spatial injury programs across mouse intracerebral haemorrhage models. **A** Spatial annotations of mouse intracerebral haemorrhage sections from collagenase-induced and autologous-blood-induced injury models. **B** Overlap between the top 500 DeSPA-ranked genes from the two injury models, identifying collagenase-specific, shared and autologous-blood-specific genes. **C** Alluvial plot showing cross-model correspondence of spatial co-expression modules among shared genes. **D** Composition of collagenase modules according to shared and model-specific gene membership. **E** GO Biological Process enrichment of collagenase spatial co-expression modules. **F** Spatial expression patterns of representative injury-associated genes, including *Pde10a, Cldn11* and *C4b*, across the collagenase trajectory.

### 2.5 DeSPA resolves developmental, regenerative and shared spatial programs in the axolotl brain

Whether brain regeneration reactivates developmental spatial programs or establishes injury-induced spatial configurations is a central question in regenerative neurobiology. We applied DeSPA to two ordered Stereo-seq series of the axolotl telencephalon: a developmental trajectory spanning stages 44, 54 and 57, juvenile, adult and metamorphic stages, and a regenerative trajectory spanning 2, 5, 10, 15, 20, 30 and 60 days post-injury [6]. The developmental and regenerative datasets comprised 36,198 and 78,123 spatial locations, respectively. Spatial annotations showed clearly resolved anatomical regions and cell-state structures in both processes, providing a basis for comparing spatial gene remodeling across development and regeneration without requiring direct coordinate alignment between the two trajectories (Fig. 5A).

Spatial co-expression module analysis was first performed separately for the developmental and regenerative trajectories using the corresponding top 500 genes ranked by DeSPA (Fig. 5B). In the developmental trajectory, the most prominent enriched module was associated with GO terms including cytoplasmic translation, macromolecule biosynthesis, translation, gene expression and protein metabolism, consistent with coordinated biosynthetic activity during brain growth and maturation. Other developmental modules were associated with synaptic and neurotransmitter-related processes, as well as extracellular matrix organization and skeletal system development, reflecting the parallel maturation of neural and structural programs. In the regenerative trajectory, the most prominent enriched GO terms similarly involved cytoplasmic translation, macromolecule biosynthesis, translation and protein metabolism (Supplementary Fig. 7). Additionally, regenerative modules were associated with pathway terms including DNA replication, RNA processing, ribosome biogenesis, neural repair, and Fc gamma receptor-mediated phagocytosis (Supplementary Fig. 8), suggesting that regeneration preserves the developmental biosynthetic backbone while recruiting injury-responsive programs. The proliferative regeneration program contained neural progenitor-associated genes, including *Sox2, Fabp7*, and *Hes5* [40–42], together with DNA-replication genes from the *Mcm2* –*Mcm7* licensing complex [43, 44]. The phagocytosis-associated program contained *Inpp5d, Dnm2*, and a *Siglec1* -like receptor, linking it to myeloid regulation and immune-associated clearance [45–47]. These patterns indicate that regeneration preserves a developmental-like biosynthetic spatial backbone but reorganizes it into additional modules linked to regeneration-specific repair programs.

We then assessed the extent to which genes prioritized during regeneration were also identified during development. Restricting the analysis to 2,697 genes detected in both datasets, we compared the top 500 genes from each process. Among these two gene sets, 246 genes were shared and 254 were process-specific (Fig. 5C). The presence of both shared and process-biased top-ranked genes is consistent with partial reuse of developmental spatial programs during regeneration, together with the recruitment of injury-associated programs that are preferentially activated during tissue repair [6].

We next compared the developmental and regenerative module assignments of the 246 shared top-ranked genes, retaining the module definitions established above. The two module systems showed moderate agreement, with the ARI (Adjusted Rand Index) of 0.476 and NMI (Normalized Mutual Information) of 0.576 (Fig. 5D), indicating that shared DeSPA genes retain partially conserved spatial co-expression organization while also being rewired between processes. The most stable correspondence was observed between development module 3 and regeneration module 5, where 37 of 40 shared genes mapped to the same regeneration module, accounting for 92.5% of the development module (Fig. 5D). At the same time, the module mapping revealed extensive regeneration-specific reorganization. Development module 1 split mainly into regeneration module 1 and module 4, whereas development module 2 mapped primarily to regeneration module 2 and development module 4 mapped predominantly to regeneration module 1 (Fig. 5D). Regeneration module composition further showed that module 5 was largely shared, with 80.3% shared genes, whereas modules 3 and 6 were dominated by regeneration-specific genes, with regeneration-specific fractions of 85.4% and 70.3%, respectively (Fig. 5E).

Finally, we linked the shared and process-specific DeSPA-prioritized gene sets to annotated cell states. Cell-state enrichment analysis revealed that shared top-ranked genes were strongly associated with cell-state signatures in both development and regeneration. For example, shared genes overlapped with vascular leptomeningeal cell (VLMC) markers in the developmental dataset (overlap = 90, FDR = 8.66 × 10**-**^28^), and in the regeneration dataset (overlap = 98, FDR = 1.62x 10 ^**-**34^). Shared genes were also enriched for choroid plexus (CP) markers in the regeneration dataset (overlap = 92, FDR = 8.85 × 10^-30^) (Fig. 5F). By contrast, regeneration-specific top-ranked genes were concentrated in regeneration-associated cell states, including regeneration intermediate progenitor cell subcluster 4 (rIPC4; overlap = 83, FDR = 1.70 × 10^-21^), regeneration intermediate progenitor cell subcluster 1 (rIPC1; overlap = 81, FDR = 2.04 × 10^-20^), reactive ependymoglial cells (reaEGC; overlap = 78, FDR = 1.34 × 10^-18^), ribosomal ependymoglial cells (ribEGC; overlap = 77, FDR = 4.45 × 10^-18^), and telencephalon neuroblasts (tlNBL; overlap = 68, FDR = 7.93 × 10^-13^) (Fig. 5F). Thus, regeneration-specific genes were preferentially associated with regenerative cell states, distinguishing them from developmental spatial-remodeling genes.

Together, these analyses show that DeSPA resolves the relationship between brain development and regeneration at the level of spatial gene programs. Development and regeneration share a conserved spatial-remodeling component enriched for translation and biosynthetic activity, while regeneration reorganizes this shared backbone into process-specific modules associated with proliferation, RNA processing, neural repair and immune-associated phagocytic activity. Thus, DeSPA distinguishes conserved developmental-like spatial programs from regeneration-specific spatial reconfiguration, providing a framework for dissecting how spatial gene regulation is reused and remodeled during tissue repair.

### 2.6 DeSPA resolves conserved and model-biased spatial injury programs across mouse ICH models

Intracerebral haemorrhage (ICH) can be induced experimentally using different strategies [48], raising the question of whether the resulting spatial programs reflect a conserved injury response or are biased by the mode of induction. We therefore applied DeSPA to the STMich mouse ICH dataset [8], which contains two ordered spatial transcriptomic series generated from collagenase-induced and autologous-blood-induced ICH models. The collagenase trajectory spanned nine time points, including naive, H3, H6, H12, D1, D3, D7, D14 and D28, whereas the autologous-blood trajectory spanned four time points, including naive, H6, D1 and D7. In the analyzed sections, the collagenase dataset contained 25,918–40,000 spatial spots per section and 23,625 genes, whereas the autologous-blood dataset contained 25,779–40,000 spatial spots per section and 23,906 genes (Fig. 6A).

DeSPA recovered a shared spatial injury program that was conserved across the two ICH models, alongside model-biased signals that distinguished the two induction strategies. Among 5,802 genes tested in both datasets, the top 500 genes ranked by DeSPA from each model comprised 337 shared genes and 163 model-specific genes (Fig. 6B). Functional enrichment analysis indicated that the shared and model-biased gene sets captured distinct aspects of the ICH response. The shared DeSPA core was enriched for behavior (FDR = 1.62 × 10^**-**13^), monoatomic ion transport (FDR = 4.11 × 10^-12^), regulation of trans-synaptic signaling (FDR = 6.23x 10^-11^), glial cell differentiation (FDR = 1.49x 10^-9^) and gliogenesis (FDR = 3.18x 10^-9^) (Supplementary Fig. 9A). These terms suggest that the conserved spatial injury program spans neuronal activity, ion homeostasis, synaptic signalling and glial remodeling. By contrast, collagenase-biased genes were enriched for synaptic transmission and plasticity-related processes, including positive regulation of synaptic transmission (FDR = 7.33 × 10^-8^), regulation of synaptic plasticity (FDR = 1.73 × 10^-7^) and long-term synaptic potentiation (FDR = 1.72 × 10 ^6^) (Supplementary Fig. 9B). Autologous-blood-biased genes were instead associated with cell junction organization (FDR = 2.87 × 10^-4^), cell-surface receptor protein tyrosine kinase signaling (FDR = 1.85 × 10^-3^), cell adhesion (FDR = 2.13 × 10^-3^) and secretion (FDR = 2.37 × 10^-3^) (Supplementary Fig. 9C). These results indicate that DeSPA separates a conserved ICH spatial injury core from model-biased programs related to synaptic plasticity, tissue organization, adhesion and secretory responses.

To determine whether the shared and model-biased genes reflect deeper organizational principles, we next examined their higher-order module architecture. Module assignments of the 337 shared genes showed an almost one-to-one correspondence between the collagenase and autologous-blood models, with the ARI of 0.906 and NMI of 0.869 (Fig. 6C). This strong module-level agreement indicates that the conserved ICH signal extends beyond shared gene identities to the spatial co-expression architecture of the injury response. Despite this prominent shared structure, each module also contained model-biased components. In the collagenase model, the fraction of shared genes within individual modules ranged from 48.4% to 85.7%, with module 4 showing the highest shared contribution (Fig. 6D). In the autologous-blood model, module-level shared fractions ranged from 52.0% to 77.2%, with modules 4 and 1 showing particularly high shared contributions of 77.2% and 74.1%, respectively (Supplementary Fig. 10). These patterns suggest that ICH spatial injury programs are organized around a stable conserved module backbone, while model-biased genes contribute additional induction-specific variation around this core.

Spatial co-expression module analysis performed separately in the two models identified biologically coherent injury-associated programs, with both shared and model-specific functional enrichments (Fig. 6E; Supplementary Fig. 11). In the collagenase model, module 1 captured a neuronal signaling program enriched for regulation of trans-synaptic signaling (FDR = 2.00 × 10^-14^), while module 2 was characterized by neurotransmitter transport and synaptic vesicle processes. Module 4 represented an oligodendrocyte- and myelin-associated program enriched for ensheathment of neurons (FDR = 4.82 × 10^**-**12^), whereas module 3 showed immune-associated functions, including positive regulation of cell killing and antigen processing and presentation. In the autologous-blood model, the neuronal programs showed similar enrichment for ion transport and trans-synaptic signaling, with module 1 showing strong enrichment for monoatomic ion transport (FDR = 9.94 × 10^-9^) and module 2 capturing synaptic organization, signal release, and neurotransmitter transport. The corresponding oligodendrocyte- and myelin-associated program, module 3, was enriched for ensheathment of neurons (FDR = 6.67 × 10^-12^) and gliogenesis (FDR = 5.54 × 10^-12^). In addition, the autologous-blood model contained a distinct injury-associated module (module 5) enriched for response to bacterium (FDR = 1.35 × 10^-4^) and response to biotic stimulus (FDR = 4.96 × 10^-4^ ), as well as antigen processing and presentation, consistent with genes such as *Vim, C4b*, and *Hba-a1*. These results reveal conserved neuronal and oligodendrocyte/myelin-associated spatial programs across the two injury models, while highlighting distinct functional enrichments related to synaptic signaling, ion transport, and immune-associated processes.

Spatial expression maps of representative genes further linked these modules to tissue-level injury patterns. The shared neuronal module was represented by *Pde10a* and *Ppp1r1b*, two striatal signaling-associated genes that were highly ranked in both ICH models [49] (Fig. 6F). A second conserved axis was marked by *Cldn11*, consistent with an oligodendrocyte and myelin-associated compartment [50] (Fig. 6F). Injury-associated modules were represented by *Vim*, together with haemorrhage- and inflammation-associated genes such as *Hba-a1* and *C4b* [51–53] (Fig. 6F). These spatial patterns connected the conserved and model-biased module structure to visible domains of neuronal dysfunction, myelin disruption, haemorrhagic lesion organization and inflammatory response.

Overall, DeSPA resolves both conserved and distinct spatial injury programs across two mouse ICH models, providing a general framework for comparative spatial transcriptomics in disease settings.

## 3 Discussion

Spatial transcriptomic atlases increasingly span diverse biological states, from developmental trajectories and disease progression to regenerative responses and therapeutic perturbations. Capturing gene expression pattern changes across these contexts is critical to elucidating developmental spatial wiring and disease-associated tissue disorganization, yet powerful tools for cross-state comparative spatial analysis remain limited. DeSPA addresses this gap by testing differential spatial patterns through within-slice, kernel-defined spatial variance components, aggregating evidence across complementary spatial kernels and slice pairs of the same state to accommodate unknown pattern geometries, and organizing top-ranked genes into cross-slice co-expression modules that capture coordinated remodeling across genes. In doing so, it transforms gene-level spatial signals into interpretable programs of tissue reorganization, providing a framework for comparative spatial transcriptomics across biological conditions.

In simulations with known ground truth, DeSPA showed robust performance across multiple forms of inter-slice heterogeneity. Across real spatial transcriptomic datasets, DeSPA recovered biologically interpretable spatial programs in distinct experimental regimes. In mouse embryogenesis, DeSPA prioritized genes with stronger stage-associated spatial remodeling, greater boundary-to-core regional redistribution and module structure corresponding to extracellular matrix organization, muscle formation, neural development and erythroid differentiation. In matched diabetic and wild-type testis sections, DeSPA produced rotation-invariant rankings, remained conservative in same-condition negative controls and resolved disease-associated programs centred on late spermatid maturation and stress-associated responses. In axolotl brain development and regeneration, DeSPA separated a shared developmental-like spatial remodeling backbone from regeneration-biased modules associated with proliferation, RNA processing, neural repair and immune-associated clearance. In mouse intracerebral haemorrhage models, DeSPA identified a conserved spatial injury core shared across collagenase-induced and autologous-blood-induced injury, while preserving model-biased programs related to synaptic plasticity, tissue organization, adhesion and inflammatory responses. These applications suggest that DeSPA is not limited to one tissue architecture or study design, but can be applied to ordered developmental trajectories, matched disease-control comparisons, regeneration series and related disease models.

DeSPA still has some limitations. First, the moment-based score-test approximation makes atlas-scale analysis feasible, but it is not equivalent to fully refitting exact likelihood-based variance-component models for every gene, slice and kernel. Approximation error may affect calibration or ranking in settings with small sample sizes, extreme sparsity, strong heteroskedasticity or highly irregular spatial sampling. Second, DeSPA reduces the need for coordinate-wise alignment, but it does not make the analysis independent of tissue geometry. Because the spatial kernels are still constructed from within-slice distance relationships, large condition-associated changes in tissue shape, missing anatomical domains, inconsistent sectioning planes or severe deformation can alter the effective geometry of the test and may be partially confounded with biological spatial remodeling. Third, the current testing framework evaluates each gene separately across slices. Although module discovery provides a downstream representation of coordinated spatial programs, the gene-level test itself does not yet borrow information across genes or integrate shared spatial covariance, pathway structure or cell-state composition during inference.

These limitations point to several extensions. Future versions of DeSPA could incorporate deformation-aware registration, multi-scale kernels or topological descriptors to improve sensitivity to complex pattern redistribution under large tissue-shape changes. Multivariate or hierarchical models could borrow information across genes and modules, improving stability for weak but coordinated spatial programs. A particularly promising direction is to extend differential spatial-pattern analysis to cell-type-specific level. Recent ct-SVG frameworks for cell-type-specific spatially variable genes (ct-SVGs) test whether a gene exhibits spatial variation within a given cell type [54, 55]. The next question is whether such cell-type-specific spatial patterns are themselves remodeled across conditions. A cell-type-aware extension of DeSPA would enable testing whether the internal spatial gene program of the same cell type is reconfigured during development, disease or regeneration, providing a more precise view of state-associated spatial regulation.

## 4 Methods

### 4.1 Identifying genes with differential spatial patterns across multiple biological conditions

DeSPA is a computational framework designed to identify genes with differential spatial patterns across multiple biological states. Each spatial transcriptomics slice is assigned a pre-specified state label, which may represent disease status, treatment, developmental stage or time point, and may contain one or more slices. The core idea of DeSPA is to model gene expression of each slice using a linear mixed model (LMM), in which kernel-based variance components quantify spatially structured variation. To robustly detect state-dependent differences in spatial patterning, DeSPA evaluates each gene by combining evidence from multiple spatial kernels within each pre-specified state comparison, and aggregating evidence across replicate slices. In addition, DeSPA provides a multi-state spatial co-expression module analysis, which groups genes with similar spatially informed co-expression patterns across states and supports downstream biological interpretation.

#### 4.1.1 Linear mixed model for gene expression

Let *M* denote the total number of slices. DeSPA takes a multi-slice spatial transcriptomics dataset in which each slice is annotated with a biological state label. The complete input is denoted by

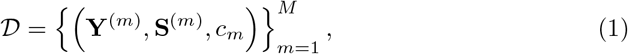

where 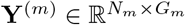 is the normalized expression matrix of slice *m*, with *N*_*m*_ spatial locations and *G*_**m**_ measured genes. Throughout, *g* denotes a gene measured in all slices involved in the relevant comparison. The corresponding spatial coordinate matrix is 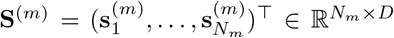 where 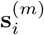 is the *D*-dimensional coordinate vector of location *i*. The label *c*_*m*_ ∈ {1,…, *C*} denotes the biological state of slice *m*, such as disease status, treatment condition, or developmental stage. Multiple slices may share the same state label, and all spatial locations within a slice inherit the label assigned to that slice.

For slice *m*, let 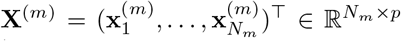 denote the full-rank design matrix, and let 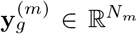 denote the expression vector of gene *g*. DeSPA uses a linear mixed model [11, 56] to separate the spatial pattern of gene expression from fixed covariate effects. Specifically, the normalized expression value at location *i* is modeled as:

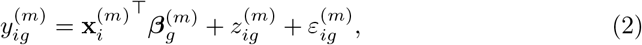

where 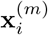 is a *p*-dimensional covariate vector that includes an intercept and *p —* 1 observed explanatory variables for slice *m* and 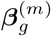 is the corresponding *p*-dimensional coefficient vector. The residual vector 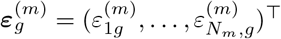 satisfies

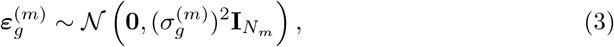

and 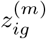 is a spatial random effect that follows a multivariate normal distribution, i.e.,

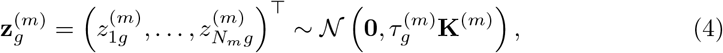

where **K**^(m)^ is an *N*_*m*_ × *N*_*m*_ kernel matrix capturing spatial correlation between locations, and 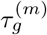 quantifies the strength of the spatial expression pattern of gene *g* in slice *m*.

Linear mixed models capture spatial information through the kernel matrix. Different kernels are sensitive to different spatial patterns: for example, a periodic kernel is naturally suited to periodic spatial structures, whereas a Gaussian kernel is more sensitive to locally concentrated patterns. DeSPA therefore uses multiple spatial kernels to robustly identify genes whose spatial patterns differ between slices. Specifically, DeSPA uses Gaussian kernel, Cauchy kernel, and periodic kernel of the form

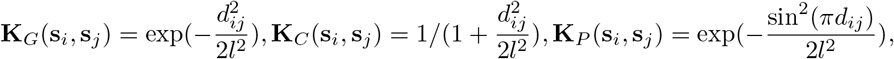

respectively, where *d*_*ij*_ = ||**s**_**i**_ — **s**_**j**_ ||_2_ is the Euclidean distance between spatial locations and *l* is the bandwidth scale. In the default setting, DeSPA uses the three kernel types above with two bandwidth scale factors, yielding six kernels per slice. Because the Euclidean distance is invariant to rotation, the resulting kernel construction is robust to rotational transformations of the spatial coordinates.

#### 4.1.2 Differential spatial pattern analysis

DeSPA estimates the slice-specific spatial variance component using a non-iterative, score-based moment approximation and then tests whether this component differs between slices. Relative to fitting every gene-wise kernel model by iterative restricted maximum likelihood (REML) [57], this approach avoids repeated variance-component optimization, while the use of multiple kernels broadens sensitivity to spatial patterns with different characteristic scales and structures.

For gene *g* in slice *m*, DeSPA first fits the fixed-effect component under the non-spatial model. The ordinary least-squares estimate of the coefficient vector is

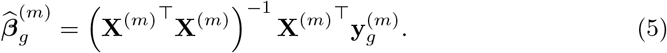

And the corresponding projection matrix onto the residual space is

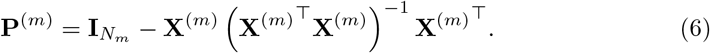

Applying this projection to the expression vector yields the residual expression vector for gene *g*:

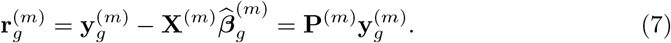

From these residuals, the non-spatial residual variance is estimated as

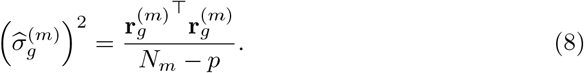

For kernel *q*, DeSPA summarizes spatial structure in the residual expression using the kernel quadratic form

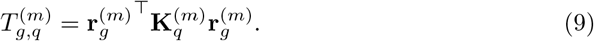

This quantity is the quadratic component underlying the variance-component score statistic. Under the LMM, its conditional expectation is

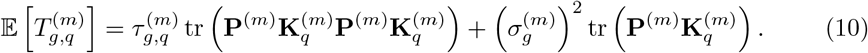

Equating the observed value of 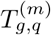 with this expectation and replacing the residual variance with its null-model estimate gives a plug-in method-of-moments estimator. Equivalently, the estimator can be viewed as a single Fisher-scoring update for the spatial variance component from the non-spatial null model [58]:

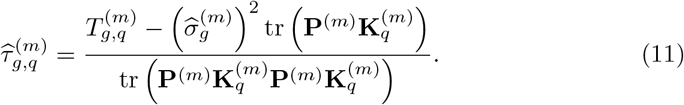

A plug-in approximation to the variance of 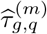 is

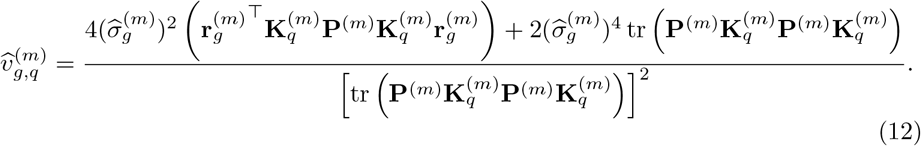

The full derivation is provided in the Supplementary Note.

DeSPA performs inference separately for each state contrast. For state *c*, let *ℳc* = *{m* : *c*_*m*_ = *c*} denote the set of slices. Consider a contrast between state *c* and *c*^0^. When each state is represented by one slice, only one cross-state slice pair is tested. When either state contains replicate slices, DeSPA evaluates all pairs in 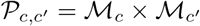. For studies with more than two states, including multiple developmental stages or time points, the same procedure is applied separately to each requested contrast, such as adjacent-time-point contrasts or all pairwise contrasts. The resulting inference is therefore specific to a state contrast rather than a single omnibus test across all states.

For slice pair 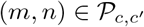 and kernel *q*, DeSPA tests

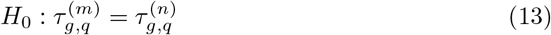

against a two-sided alternative. Treating slices as independent sampling units, the corresponding standardized statistic is

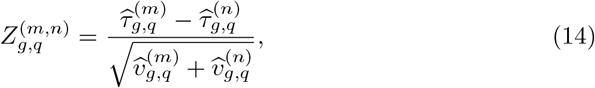

and the kernel-specific two-sided *p*-value is

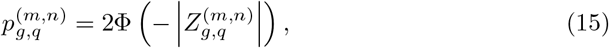

where Φ (·) is the standard normal cumulative distribution function.

DeSPA combines the resulting *p*-values in two stages. First, for a fixed gene and slice pair, the kernel-specific *p*-values are combined into a slice-pair *p*-value, denoted by 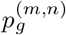, using the aggregated Cauchy association test (ACAT) [19, 20]. This step integrates evidence from kernels with different sensitivities while accommodating their correlation. Second, for a state contrast (*c, c*’), DeSPA combines the slice-pair *p*-values 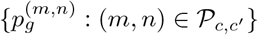 using Fisher’s method to obtain a contrast-specific gene-level *p*-value. Within each state contrast, the resulting gene-level *p*-values are adjusted using the Benjamini–Hochberg procedure [21]. By default, DeSPA evaluates all unordered pairs of distinct state labels. Users may instead provide pre-specified state contrasts or slice pairs for targeted analyses.

#### 4.1.3 Cross-slice spatial co-expression module analysis

Following differential spatial pattern analysis, DeSPA utilizes the top-ranked genes to identify spatial co-expression modules across slices. This module identification proceeds in two steps: DeSPA first infers slice-specific gene co-expression relationships via a spatial smoothing operator, and subsequently integrates these local networks into unified cross-slice modules.

Specifically, for a gene set *G***\*** and slice *m*, DeSPA extracts the standardized expression matrix 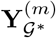 to construct a slice-specific gene network. To mitigate spatial noise, DeSPA applies a spatial *k*-nearest-neighbour smoothing operator **W**^(*m*)^ (by default, *k* = 10) to the expression profiles:

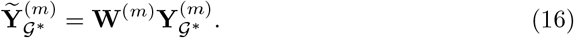

For genes *g,h* ∈ *G\**, their co-expression strength in slice *m* is defined as the nonnegative Pearson correlation

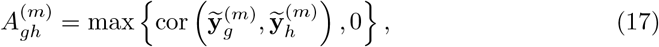

where 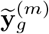 is the column of 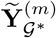corresponding to gene *g*. The slice-specific adjacency matrix is denoted by 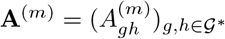.

To obtain spatial co-expression modules across slices, DeSPA treats 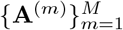 as layers of a multiplex gene network with the common vertex set G^*^. Let *u*_g_ denote the module assignment of gene g, and let 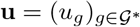 denote the shared assignment vector. DeSPA estimates u by maximizing the layer-summed modularity objective [59, 60]:

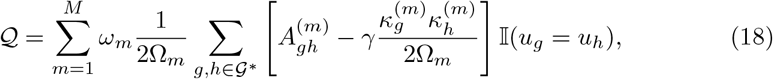

where 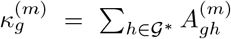 is the weighted degree of gene *g* in layer 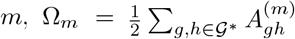 is the total edge weight of that layer, *7* is the resolution parameter, *ω*_*m*_ is the layer weight and *I*(·) is the indicator function. In the default setting, all layers are assigned equal weight. DeSPA optimizes this objective using multiplex Leiden community detection [61]. This produces one module label for each gene, while allowing the edge weights supporting that module structure to vary by slice. The resulting modules therefore represent cross-slice spatial co-expression modules that can be compared across biological conditions without requiring subsequent alignment of independently clustered slice-specific modules.

#### 4.1.4 Computational complexity

The primary computational bottleneck in traditional variance-component models lies in iterative restricted maximum likelihood (REML) fitting, which typically requires inversion or eigen-decomposition of dense covariance matrices. For a slice with *Nb* spatial locations, this operation has cubic complexity, 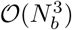, for each dense covariance calculation. When spatial patterns are compared across many genes, slices and kernels, repeating this operation for every gene-kernel combination becomes computationally intractable. DeSPA circumvents this bottleneck by reducing spatial variance estimation to sparse matrix operations.

Let *N*_*b*_ denote the number of spatial locations in slice *b, Q* the number of spatial kernels, *G* the number of common genes tested and *k* the number of spatial neighbours retained in the sparse kernel representation. The complexity of DeSPA is determined by two main steps. First, DeSPA constructs sparse spatial kernels from the tissue coordinates. Using a tree-based nearest-neighbour search, this step has an approximate complexity of *O*(*N*_*b*_ log *N*_*b*_), and evaluating the *Q* kernels on the retained neighbour graph takes *O(QkN*_*b*_). Second, instead of performing 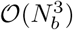 REML fitting, DeSPA estimates the spatial variance components using sparse matrix-vector operations involving the residual expression vector and the kernel matrix. This moment-based approximation reduces the dominant per-slice complexity to *O*(*QkN*_*b*_ *G*). Consequently, the total computational cost across all *B* slices is approximately bounded by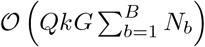 . Because *Q* and *k* are fixed by default, DeSPA scales approximately linearly with the number of tested genes and the total number of spatial locations. This acceleration enables systematic differential spatial-pattern analysis across multi-slice spatial atlases without repeated dense REML optimization.

### 4.2 Compared methods

To evaluate DeSPA, we benchmarked it against River and two Moran’s *I* -based rankings.

**River** [18] is an interpretable deep-learning framework for prioritizing genes with perturbation-responsive spatial patterns. For each simulated dataset, River was run on the corresponding paired-slice input using its default parameters. Genes were then ranked in descending order according to the River score. The resulting ranked list was evaluated using the same power and empirical FDR procedure as DeSPA.

**Moran’s** *I* [62] was used as a spatial-autocorrelation baseline. For each gene, Moran’s *I* was computed separately in the two simulated slices, denoted by 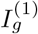and 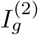. We considered two Moran’s *I*-based scores. The absolute-change score was defined as

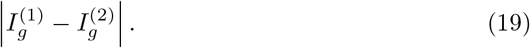

The relative-change score was defined as

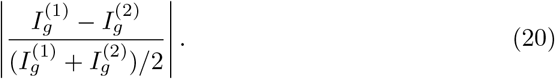

For both Moran’s *I* baselines, genes were ranked in descending order according to the corresponding score. The resulting ranked lists were evaluated using the same power and empirical FDR procedure as DeSPA.

### 4.3 Datasets

#### 4.3.1 Simulation datasets

To systematically benchmark the methods for differential spatial-pattern analysis, we generated paired slices under different conditions with known ground-truth DSEPs. The simulation design separated slice-level heterogeneity from gene-level spatial-pattern changes. The three slice-level scenarios described global differences between the two slices, including unequal sampling density of spatial locations, broad nuisance fixed effects and geometric deformation. The gene-level settings then determined whether and how the underlying spatial pattern of each gene changed between slices. Each paired-slice benchmark comprised 200 genes, including 100 null genes, 50 interference genes and 50 positive genes. Null and interference genes were treated as negative controls: null genes had no target spatial-pattern change, whereas interference genes exhibited slice-to-slice expression perturbations that did not alter their underlying spatial pattern. Positive genes represented genuine differential spatial-pattern genes. This design allowed us to evaluate the power of each method to detect ground-truth differential spatial patterns and its robustness to non-target between-slice variation.

The three slice-level scenarios were set as follows.

In scenario 1, both slices were sampled uniformly from the same square tissue region, [0,10] x [0,10]. Slice 1 contained 300 spatial locations, whereas slice 2 contained 150 spatial locations. The tissue region and coordinate geometry were therefore held constant, and only the sampling density of spatial locations differed between slices. This setting modelled unequal spatial sampling density between paired slices.

In scenario 2, both slices contained 300 spatial locations sampled from the same square tissue region. Slice 2 additionally included a broad binary fixed-effect covariate that partitioned the locations into an affected region and a baseline region. This covariate introduced slice-specific nuisance variation, representing, for example, platform-, tissue-region- or batch-associated effects, but was not used to determine the ground truth DSEP status of genes. This setting modelled broad slice-specific confounding while preserving identical spatial sampling between the paired slices.

In scenario 3, gene expression patterns were generated in a shared rectangular reference coordinate system, with *x* ∈ [0, 5] and *y* ∈ [0,10]. The observed coordinates of slice 1 were identical to the reference coordinates, whereas the observed physical coordinates of slice 2 were transformed using the Cartesian-to-polar bending map

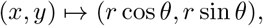

where *r* = 5 + *x* and θ= ((*y/*10) — 0.5) *π/*2. Gene-level spatial patterns were defined in the shared reference coordinate system before the transformation was applied. This setting modelled geometric warping of slice 2 coordinates while preserving a common reference system for generating the underlying expression patterns.

Across all three scenarios, expression at each spatial location was generated from a baseline mean of 2.0 with additive Gaussian residual noise of standard deviation 0.5. Gene-specific spatial signals and, where applicable, slice-specific nuisance effects were then added according to the gene category and slice-level scenario. Spatial signals were constructed from three smooth prototype patterns defined over the reference coordinate system: a ring and a Gaussian blob, both centred at the spatial centroid, and a vertical Gaussian strip centred at the mean reference *x* coordinate. Unless stated otherwise, the spatial-signal amplitude was set to 2.0.

Null genes contained no gene-specific spatial signal in either slice and were therefore negative for DSEPs. In scenario 2, the fixed-effect nuisance term was additionally applied to their expression values in slice 2. Because this term represented slice-specific nuisance variation rather than a change in the ground-truth gene-specific spatial pattern, null genes remained negative controls for evaluating false discoveries.

Interference genes were included to assess whether methods incorrectly interpreted non-target expression changes as differential spatial-pattern signals. These genes shared the same underlying strip-shaped spatial pattern across slices and were also negative for DSEPs. Within each benchmark, all 50 interference genes were subjected to one of two perturbations in slice 2. Mean-shift interference was generated by adding 5.0 to the spatially uniform baseline mean, whereas noise-inflation interference was generated by increasing the residual-noise standard deviation from 0.5 to 1.0. In scenario 2, the fixed-effect contribution was also added to interference-gene expression in slice 2. These perturbations altered the overall expression level or residual variability without changing the underlying spatial pattern, and interference genes therefore remained negative controls.

Positive genes represented genuine changes in gene-specific spatial patterns. Within each benchmark, all 50 positive genes were generated under one of five settings. Two topology-change settings comprised ring-to-blob and blob-to-strip transitions. Two pattern-emergence settings comprised blob emergence and strip emergence, in which slice 1 contained no gene-specific spatial signal and slice 2 contained the corresponding spatial pattern. In the intensity-change setting, both slices contained the same blob pattern, but its amplitude increased from 2.0 in slice 1 to 4.0 in slice 2.

The full factorial combination of three slice-level scenarios, two interference settings and five positive settings yielded 30 (3 × 2 × 5) paired-slice benchmark configurations, including 10 configurations for each slice-level scenario. The simulated expression matrices and observed spatial coordinates were retained for downstream analysis, together with the fixed-effect covariate in scenario 2. Positive genes defined the ground truth for power evaluation, whereas null and interference genes were used to assess false-discovery control.

#### 4.3.2 Real datasets

We applied DeSPA to four publicly available spatial transcriptomics datasets spanning embryonic development, metabolic disease, regeneration and brain injury.

##### Stereo-seq mouse embryo development data

This dataset [5] profiles mouse embryo development using the Stereo-seq platform. Our analysis used eight developmental stages, E9.5, E10.5, E11.5, E12.5, E13.5, E14.5, E15.5 and E16.5. The analyzed slices contained 5,913–121,767 spatial locations before preprocessing. After applying the common filtering and gene-intersection procedure described below, the retained gene set was used for stage-wise DeSPA ranking and comparative analyses. Tissue annotations provided with the dataset were used for spatial visualization and local region quantification.

##### Slide-seq mouse diabetes testis data

This dataset [7] contains mouse testis spatial transcriptomics profiles generated using Slide-seq. It is composed of six Slide-seq slices from three leptin-deficient diabetic mice and three matched wild-type mice. The individual slices contained 27,194–42,776 spatial locations and 23,514–24,450 measured genes before preprocessing. DeSPA was applied to the common filtered gene set across matched samples. Cell-type annotations were mapped to nine testis cell populations, including spermatid, spermatocyte, spermatogonium, Sertoli, Leydig, endothelial and macrophage populations, and were used for tissue visualization.

##### Stereo-seq axolotl brain regeneration data

This dataset [6] profiles axolotl telencephalon development and brain regeneration using the Stereo-seq platform. We analyzed two related processes. The developmental process included one analysed sample from each of six developmental stages: Stage44, Stage54, Stage57, juvenile/control, adult and metamorphosis. The regeneration process included an uninjured control and one analyzed sample from each injured time point at 2, 5, 10, 15, 20, 30 and 60 days post-injury. Across these analyzed samples, the developmental and regeneration processes contained 36,198 and 78,123 spatial locations, respectively, before process-specific filtering. These data were used to compare top-ranked genes and shared spatial co-expression modules between developmental and regenerative processes.

##### Stereo-seq mouse intracerebral haemorrhage data

This dataset [8] profiles mouse intracerebral haemorrhage (ICH) using the Stereo-seq platform. We analysed two ICH models: collagenase-induced ICH and autologous-blood ICH. The full collagenase and autologous-blood datasets contained 3,471,721 and 816,207 spatial locations, respectively, before slice selection. For the collagenase model, DeSPA was applied to one selected slice from each of nine time points: Naive, H3, H6, H12, D1, D3, D7, D14 and D28. For the autologous-blood model, we used one selected slice from Naive, H6, D1 and D7 as a model-comparison reference. The selected collagenase and autologous-blood slices contained 711,045 and 264,062 spatial locations, respectively. Region and cell-type annotations provided in the dataset were used for spatial visualization, region-level quantification and gene-set score analysis.

### 4.4 Details of analyses

#### 4.4.1 Data preprocessing

For real spatial transcriptomics datasets, preprocessing was performed using `Scanpy` [63] to obtain comparable expression matrices across slices within each analysis. For the mouse diabetes testis and Stereo-seq axolotl brain regeneration datasets, genes detected in fewer than 50 spatial locations and locations with fewer than 10 detected genes were removed. For the higher-resolution Stereo-seq mouse embryo development and Stereo-seq mouse ICH datasets, genes detected in fewer than 500 spatial locations and locations with fewer than 10 detected genes were removed. Each raw-count slice was subsequently library-size normalized to a total count of 10,000 per location and log-transformed using `log1p`.

After slice-level preprocessing, genes were intersected across all slices included in the corresponding comparison, and only this common filtered gene set was used for DeSPA ranking and downstream analyses. This ensured that differential spatial pattern testing, spatial pattern remodeling analysis, co-expression module analysis and functional enrichment were all performed on the same genes within each dataset.

#### 4.4.2 Evaluation on simulation data

Simulation performance was evaluated against the known positive gene labels assigned during data generation. For each simulated dataset, the reference positive set contained 50 genes with true differential spatial patterns, whereas null and interference genes were treated as non-positive for empirical error estimation.

We compared DeSPA with River, Moran’s *I* -based ranking with absolute-change and relative-change score. For each method, we evaluated the full ranked list using two metrics: statistical power and empirical false discovery rate (FDR). At each rank cutoff, corresponding to the selected top-ranked genes up to that point, we counted the cumulative numbers of positive genes and non-positive genes, denoted by TP and FP, respectively. Power was defined as

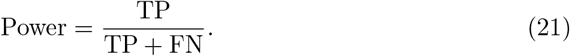

Empirical FDR was defined as

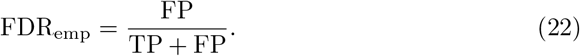

Power (sensitivity) measured the fraction of true differential spatial-pattern genes recovered by a method, whereas empirical FDR measured the proportion of non-positive genes among all selected genes at the same rank cutoff. For each target FDR threshold, we reported the power at the deepest rank cutoff whose empirical FDR did not exceed that threshold. Target FDR thresholds were set to 0.01, 0.03, 0.05, 0.075 and 0.10. Evaluation curves were generated separately for each spatial scenario and interference setting. The primary scalar summary used in the Results was power at a target empirical FDR 0.05.

#### 4.4.3 Evaluation on real data

##### Spatial autocorrelation difference analysis across developmental stages

To evaluate whether top-ranked genes captured developmental changes in global spatial autocorrelation, we used Moran’s *I* [62], a standard measure of spatial autocorrelation, to summarize the spatial organization of each gene in each stage. Specifically, for ordered developmental stages, let *I*_**g**,**t**_ denote the Moran’s *I* for gene *g* at stage *t*, computed on a spatial *k*-nearest-neighbour (*k*-NN) graph. The *k*-NN graphs were constructed with *k* = 15. To mitigate the computational overhead imposed by large slices, we downsampled each stage to a maximum of 40,000 spatial locations. The spatial autocorrelation difference score, 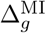, was defined as the mean absolute change in Moran’s *I* between adjacent stages:

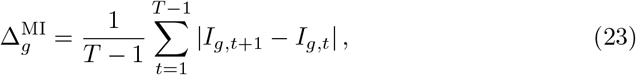

where *T* represents the total number of ordered stages. We utilized this metric to benchmark DeSPA against the comparison methods. To ensure cutoff robustness, we evaluated the mean 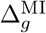 among the top 50, 100, 200, 500 and 1,000 genes from each ranking. Statistical differences between top-500 gene sets were assessed via two-sided Wilcoxon rank-sum tests.

##### Local spatial pattern difference analysis across developmental stages

Spatial autocorrelation summarizes global spatial organization but does not specify where spatial-pattern differences occur within the tissue. To quantify local spatial pattern differences across developmental stages, we measured the redistribution of gene expression across boundary-to-core tissue regions. Specifically, we calculated the distance from each spatial location to the tissue boundary and normalized this distance within each stage, providing comparable tissue regions across stages without requiring pre-defined anatomical annotations. Locations with normalized distance *δ*_*i*_ ≤ 0.30 were assigned to the boundary region, locations with *δ*_*i*_ ≥ 0.70 were assigned to the core region, and the remaining locations were assigned to the transition region.

Let *ℛ* = {boundary, transition, core} denote the set of three regions defined above. For gene *g*, region *u* ∈ ℛ, and stage *t*, we computed the regional expression fraction:

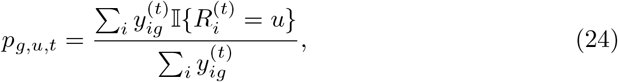

where 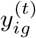 denotes the expression value of gene *g* at spatial location *i* in stage *t*, and 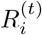 denotes the boundary-to-core region label of location *i* in stage *t*. This fraction measures the contribution of region *u* to the total expression of gene *g* at stage *t*.

We then summarized local spatial pattern differences for each gene using two metrics. The first metric, the all-stage local spatial pattern difference score, was defined as

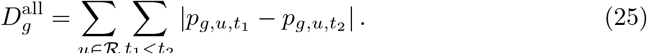

The second metric, the adjacent-stage local spatial pattern difference score, was defined as

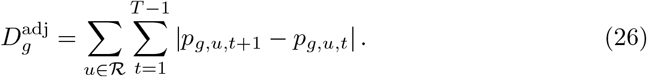

The all-stage local spatial pattern difference score summarizes redistribution across all pairs of developmental stages, whereas the adjacent-stage local spatial pattern difference score focuses on redistribution between consecutive stages. Higher values indicate stronger local spatial pattern differences across boundary, transition and core regions during development. We compared the distributions of these metrics among DeSPA, River and HVG top-ranked gene sets using the same top-500 cutoff and Wilcoxon rank sum testing framework. For visualization and reporting, both local spatial pattern difference scores were multiplied by 100.

##### Rotation robustness analysis

Spatial transcriptomics samples from different states are rarely acquired in identical orientations; therefore, a robust differential spatial-pattern method should be invariant to the rigid rotation of any individual sample. To evaluate this property, we utilized the matched diabetic and wildtype testis slices and rotated the spatial coordinates of the diabetic slices by θ ∈ {45**°**, 90**°**, 135**°**, 180**°**, 225**°**, 270**°**, 315**°**}. For each rotation angle θ, DeSPA was rerun to obtain a ranked gene list *π*_*θ*_, which was compared with the ranking *π*_θ_ obtained from the original coordinates. We quantified top-ranked gene stability using the Jaccard index of the top *K* = 500 genes:

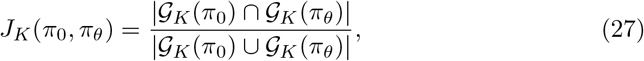

where *G*_*K*_(*π*) denotes the top *K* genes in ranking *π*. An ideally robust method is expected to yield identical rankings (*J*_*K*_ = 1) regardless of coordinate transformations. Furthermore, we assessed genome-wide ranking consistency using the Spearman rank correlation across the common gene universe, and visually compared the 0°reference ranking against the 90*°* rotation. The identical evaluation framework was applied to River as a control method.

##### Cross-process comparison of DSEP genes and spatial co-expression modules

To investigate the top-ranked genes and co-expression modules underlying spatial pattern remodeling across related biological processes, we analysed the axolotl development and regeneration datasets separately and compared their top-ranked genes and module structures. Specifically, DeSPA was first applied within each process to rank genes based on their differential spatial patterning across ordered stages. The top 500 ranked genes from the two processes were subsequently partitioned into shared, development-specific, and regeneration-specific gene sets. To compare their modular organization, we performed spatial co-expression module analysis separately for development and regeneration using the top 500 genes from each process, with 10 spatial neighbours, top-30 retained gene-gene edges and resolution parameter *γ* = 1.0. For genes that were top-ranked in both processes, we tabulated their development and regeneration module assignments and visualized the resulting module correspondences using an alluvial diagram. Finally, for each regeneration module, we further calculated the fraction of genes belonging to the shared and regeneration-specific categories, thereby distinguishing conserved module components from regeneration-biased components.

##### Cell-state enrichment analysis of DSEP genes

To determine which annotated cell states were associated with process-specific DSEP genes, we performed an enrichment analysis for the axolotl regeneration dataset. The cell-state labels were extracted from the provided annotations. Let S_**c**_ denote a query gene set, where *c* represents the shared, development-specific, or regeneration-specific top-500 category. For each annotated state *s* and considered gene *g*, we computed a state-specific relative expression score:

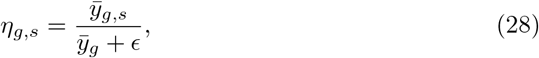

where 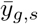 is the mean log-normalized expression of gene *g* in locations annotated as state 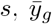 is its mean log-normalized expression across all analyzed locations, and *ε* = 10^-8^ avoids division by zero. The top 300 genes ranked by *y*_*g*_,*s* were designated as the state signature gene set ℳ_**s**_ . Enrichment of each query set *S*_**c**_ in each signature set ℳ_***s***_ was quantified by the overlap *k*_***c***,***s***_ = |*S*_***c***_ ηℳ_**s**_ | and evaluated using a one-sided hypergeometric test [64]. The resulting *p*-values were corrected using the Benjamini– Hochberg procedure. In parallel, we computed a spatial gene-set score for each location as the mean log-normalized expression of genes in S_**c**_ and summarized this score within each annotated state by its mean value. Mean gene-set scores were further standardized across states for visualization.

##### Functional enrichment analysis of DSEP genes

We performed functional enrichment analysis to characterize the biological processes over-represented within selected spatial gene sets. To avoid background bias, the corresponding tested gene universe for each dataset was utilized as the statistical background. Enrichment was assessed using a one-sided hypergeometric over-representation test [64], implemented in Python with `scipy.stats.hypergeom`, and *p*-values were adjusted using the Benjamini–Hochberg procedure. For murine datasets, we utilized the MSigDB Mouse Gene Ontology (GO) Biological Process collection (v2025.1). For the axolotl development and regeneration dataset, queries were mapped to human orthologs and evaluated against the 2025 GO Biological Process database. For axolotl regeneration modules, Kyoto Encyclopedia of Genes and Genomes (KEGG) pathway enrichment was additionally performed using the KEGG 2021 Human gene-set library. Functional terms with a false discovery rate (FDR) *<* 0.05 were considered statistically significant.

## >4.5 Code availability

The DeSPA software code is publicly available at https://github.com/kotone-429/DeSPA.

## 4.6 Data availability

The data analyzed in this study can be accessed through: Stereo-seq mouse embryo development dataset: CNGB database under accession code CNP0001543; Slide-seq mouse diabetes dataset: https://www.dropbox.com/scl/fi/dr10k6x92hxdnuijgdobn/Testis_Slideseq_Data.zip?rlkey=yy5odlkqqynfped6rixwcnk5l&e=1&dl=0; Stereo-seq axolotl brain regeneration dataset: CNGB database under accession code CNP0002068; Stereo-seq mouse ICH dataset: Spatial Transcript Omics Database (STOmics DB) under accession code `STT0000047`.

## Author contributions

Conceptualization: SZ and QS; methodology: HW; software, HW; validation: HW, QS; formal analysis: HW; investigation: HW; data curation: HW; writing—original draft preparation: HW; writing—review and editing: all authors; visualization: HW; supervision: SZ and ZY; project administration: HW; funding acquisition, SZ. All authors have read and agreed to the published version of the manuscript.

