## Supplementary materials for "DeSPA uncovers genes with cross-state spatial pattern remodeling and coordinated spatial programs across multi-slice spatial transcriptomics atlases"

### Supplementary Information for "DeSPA uncovers genes with differential spatial patterns across biological states"

#### Supplementary Note 1: Derivation of the Score-Based Moment Estimator for Differential Spatial-Pattern Analysis

The main Methods section describes the non-iterative estimator used by DeSPA to quantify the kernel-specific spatial variance component within each slice. Here, we provide the intermediate steps connecting the residualized linear mixed model, the quadratic-form moment equation, the one-step Fisher-scoring estimator, and its plug-in variance approximation. Throughout this note, we consider a fixed gene $g$, slice $m$, and spatial kernel $q$, and retain the indices used in the main text. All expectations and variances are conditional on the fixed design matrix $\mathbf{X}^{(m)}$.

##### Residualized linear mixed model

For kernel $q$, the vector form of the linear mixed model is

$$\mathbf{y}_g^{(m)} = \mathbf{X}^{(m)}\boldsymbol{\beta}_g^{(m)} + \mathbf{z}_g^{(m)} + \boldsymbol{\varepsilon}_g^{(m)}, \quad \mathbf{z}_g^{(m)} \sim \mathcal{N}\left(\mathbf{0}, \tau_{g,q}^{(m)}\mathbf{K}_q^{(m)}\right), \quad \boldsymbol{\varepsilon}_g^{(m)} \sim \mathcal{N}\left(\mathbf{0}, (\sigma_g^{(m)})^2\mathbf{I}_{N_m}\right), \tag{S1}$$

where the spatial random effect and residual noise are independent. Let

$$\mathbf{P}^{(m)} = \mathbf{I}_{N_m} - \mathbf{X}^{(m)}\left(\mathbf{X}^{(m)\top}\mathbf{X}^{(m)}\right)^{-1}\mathbf{X}^{(m)\top} \tag{S2}$$

be the ordinary least-squares residual projection matrix. Because $\mathbf{X}^{(m)}$ has full column rank, $\mathbf{P}^{(m)}$ is symmetric and idempotent, satisfies $\mathbf{P}^{(m)}\mathbf{X}^{(m)} = \mathbf{0}$, and has trace $\mathrm{tr}(\mathbf{P}^{(m)}) = N_m - p$. The residualized expression vector is therefore

$$\mathbf{r}_g^{(m)} = \mathbf{P}^{(m)}\mathbf{y}_g^{(m)} = \mathbf{P}^{(m)}\mathbf{z}_g^{(m)} + \mathbf{P}^{(m)}\boldsymbol{\varepsilon}_g^{(m)}. \tag{S3}$$

After integrating over the spatial random effect and residual noise, its covariance is

$$\begin{aligned}\boldsymbol{\Sigma}_{r,g,q}^{(m)} &= \mathrm{Cov}\left(\mathbf{r}_g^{(m)}\right) \\ &= \mathbf{P}^{(m)}\left\{\tau_{g,q}^{(m)}\mathbf{K}_q^{(m)} + (\sigma_g^{(m)})^2\mathbf{I}_{N_m}\right\}\mathbf{P}^{(m)} \\ &= \tau_{g,q}^{(m)}\mathbf{P}^{(m)}\mathbf{K}_q^{(m)}\mathbf{P}^{(m)} + (\sigma_g^{(m)})^2\mathbf{P}^{(m)}.\end{aligned} \tag{S4}$$

Under the non-spatial working model, $\tau_{g,q}^{(m)} = 0$, and

$$\mathbb{E}\left[\mathbf{r}_g^{(m)\top}\mathbf{r}_g^{(m)}\right] = (\sigma_g^{(m)})^2\,\mathrm{tr}\left(\mathbf{P}^{(m)}\right) = (\sigma_g^{(m)})^2(N_m - p). \tag{S5}$$

This gives the null-model residual-variance estimator used in the main text,

$$\left(\widehat{\sigma}_g^{(m)}\right)^2 = \frac{\mathbf{r}_g^{(m)\top}\mathbf{r}_g^{(m)}}{N_m - p}. \tag{S6}$$

#### Quadratic-form moment equation

For kernel $q$, DeSPA summarizes the residual spatial structure using

$$T_{g,q}^{(m)} = \mathbf{r}_g^{(m)\top}\mathbf{K}_q^{(m)}\mathbf{r}_g^{(m)}. \tag{S7}$$

For a zero-mean random vector $\mathbf{a}$ with covariance $\boldsymbol{\Sigma}$ and a symmetric matrix $\mathbf{A}$, $\mathbb{E}(\mathbf{a}^\top\mathbf{A}\mathbf{a}) = \operatorname{tr}(\mathbf{A}\boldsymbol{\Sigma})$. Applying this identity to Eq. (S7) and using Eq. (S4) gives

$$\begin{aligned}\mathbb{E}\Big[T_{g,q}^{(m)}\Big] &= \operatorname{tr}\Big(\mathbf{K}_q^{(m)}\boldsymbol{\Sigma}_{r,g,q}^{(m)}\Big)\\ &= \tau_{g,q}^{(m)}\operatorname{tr}\Big(\mathbf{P}^{(m)}\mathbf{K}_q^{(m)}\mathbf{P}^{(m)}\mathbf{K}_q^{(m)}\Big) + (\sigma_g^{(m)})^2\operatorname{tr}\Big(\mathbf{P}^{(m)}\mathbf{K}_q^{(m)}\Big).\end{aligned} \tag{S8}$$

where cyclic invariance of the trace and the symmetry of $\mathbf{P}^{(m)}$ and $\mathbf{K}_q^{(m)}$ were used. Define the kernel-specific constants

$$A_q^{(m)} = \operatorname{tr}\Big(\mathbf{P}^{(m)}\mathbf{K}_q^{(m)}\mathbf{P}^{(m)}\mathbf{K}_q^{(m)}\Big), \qquad B_q^{(m)} = \operatorname{tr}\Big(\mathbf{P}^{(m)}\mathbf{K}_q^{(m)}\Big). \tag{S9}$$

Because $\mathbf{P}^{(m)}\mathbf{K}_q^{(m)}\mathbf{P}^{(m)}$ is symmetric, $A_q^{(m)} = \operatorname{tr}\{(\mathbf{P}^{(m)}\mathbf{K}_q^{(m)}\mathbf{P}^{(m)})^2\} = \|\mathbf{P}^{(m)}\mathbf{K}_q^{(m)}\mathbf{P}^{(m)}\|_F^2 \geq 0$, and it is positive for a non-degenerate kernel after removal of the fixed-effect subspace. Equation (S8) can then be written as

$$\mathbb{E}\Big[T_{g,q}^{(m)} - (\sigma_g^{(m)})^2 B_q^{(m)}\Big] = \tau_{g,q}^{(m)} A_q^{(m)}. \tag{S10}$$

Equating the observed quadratic form with its model expectation and replacing the residual variance by Eq. (S6) yields

$$\widehat{\tau}_{g,q}^{(m)} = \frac{T_{g,q}^{(m)} - \left(\widehat{\sigma}_g^{(m)}\right)^2 B_q^{(m)}}{A_q^{(m)}} = \frac{T_{g,q}^{(m)} - \left(\widehat{\sigma}_g^{(m)}\right)^2 \operatorname{tr}\Big(\mathbf{P}^{(m)}\mathbf{K}_q^{(m)}\Big)}{\operatorname{tr}\Big(\mathbf{P}^{(m)}\mathbf{K}_q^{(m)}\mathbf{P}^{(m)}\mathbf{K}_q^{(m)}\Big)}. \tag{S11}$$

Thus, the estimator is a method-of-moments estimate obtained after removing the fixed-effect subspace and subtracting the expected contribution of non-spatial residual noise from the kernel quadratic form.

#### Equivalent one-step Fisher-scoring representation

The same estimator can be obtained from the restricted-likelihood score. For the fixed gene, slice, and kernel, let

$$\mathbf{V}_{g,q}^{(m)}(\tau) = \tau\mathbf{K}_q^{(m)} + (\sigma_g^{(m)})^2\mathbf{I}_{N_m}, \tag{S12}$$

and define the covariance-weighted residual projector

$$\mathbf{P}_V^{(m)}(\tau) = \mathbf{V}_{g,q}^{(m)}(\tau)^{-1} - \mathbf{V}_{g,q}^{(m)}(\tau)^{-1}\mathbf{X}^{(m)}\left\{\mathbf{X}^{(m)\top}\mathbf{V}_{g,q}^{(m)}(\tau)^{-1}\mathbf{X}^{(m)}\right\}^{-1}\mathbf{X}^{(m)\top}\mathbf{V}_{g,q}^{(m)}(\tau)^{-1}. \tag{S13}$$

With $(\sigma_g^{(m)})^2$ held at its null-model estimate, the restricted-likelihood score for the spatial variance component is

$$U_{g,q}^{(m)}(\tau) = \frac{1}{2}\left\{\mathbf{y}_g^{(m)\top}\mathbf{P}_V^{(m)}(\tau)\mathbf{K}_q^{(m)}\mathbf{P}_V^{(m)}(\tau)\mathbf{y}_g^{(m)} - \operatorname{tr}\Big[\mathbf{P}_V^{(m)}(\tau)\mathbf{K}_q^{(m)}\Big]\right\}. \tag{S14}$$

At the non-spatial null, $\tau = 0$, we have $\mathbf{V}_{g,q}^{(m)}(0) = (\sigma_g^{(m)})^2\mathbf{I}_{N_m}$ and $\mathbf{P}_V^{(m)}(0) = (\sigma_g^{(m)})^{-2}\mathbf{P}^{(m)}$. Therefore,

$$U_{g,q}^{(m)}(0) = \frac{T_{g,q}^{(m)} - (\sigma_g^{(m)})^2 B_q^{(m)}}{2(\sigma_g^{(m)})^4}. \tag{S15}$$

The corresponding expected Fisher information is

$$\mathcal{I}_{g,q}^{(m)}(0) = \frac{1}{2}\operatorname{tr}\left[\mathbf{P}_V^{(m)}(0)\mathbf{K}_q^{(m)}\mathbf{P}_V^{(m)}(0)\mathbf{K}_q^{(m)}\right] = \frac{A_q^{(m)}}{2(\sigma_g^{(m)})^4}. \tag{S16}$$

Starting from $\tau = 0$, one Fisher-scoring update is

$$\tau^{(1)} = 0 + \left\{\mathcal{I}_{g,q}^{(m)}(0)\right\}^{-1} U_{g,q}^{(m)}(0) = \frac{T_{g,q}^{(m)} - (\sigma_g^{(m)})^2 B_q^{(m)}}{A_q^{(m)}}. \tag{S17}$$

Replacing $(\sigma_g^{(m)})^2$ with the null-model estimate in Eq. (S6) recovers Eq. (S11). This establishes the equivalence between the moment estimator and a single Fisher-scoring update from the non-spatial model, without iterative variance-component optimization.

#### Plug-in variance approximation and between-slice statistic

To obtain the variance approximation used by DeSPA, write the residual vector conditional on the realized spatial random effect as

$$\mathbf{r}_g^{(m)} = \boldsymbol{\mu}_g^{(m)} + \mathbf{e}_g^{(m)}, \qquad \boldsymbol{\mu}_g^{(m)} = \mathbf{P}^{(m)}\mathbf{z}_g^{(m)}, \qquad \mathbf{e}_g^{(m)} = \mathbf{P}^{(m)}\boldsymbol{\varepsilon}_g^{(m)} \sim \mathcal{N}\Big(\mathbf{0}, (\sigma_g^{(m)})^2\mathbf{P}^{(m)}\Big). \tag{S18}$$

For a Gaussian vector $\mathbf{a} \sim \mathcal{N}(\boldsymbol{\mu}, \boldsymbol{\Sigma})$ and a symmetric matrix $\mathbf{A}$,

$$\operatorname{Var}(\mathbf{a}^\top\mathbf{A}\mathbf{a}) = 2\operatorname{tr}(\mathbf{A}\boldsymbol{\Sigma}\mathbf{A}\boldsymbol{\Sigma}) + 4\boldsymbol{\mu}^\top\mathbf{A}\boldsymbol{\Sigma}\mathbf{A}\boldsymbol{\mu}. \tag{S19}$$

Applying Eq. (S19) to $T_{g,q}^{(m)}$ conditional on $\mathbf{z}_g^{(m)}$ gives

$$\begin{aligned}\operatorname{Var}\Big(T_{g,q}^{(m)} \mid \mathbf{z}_g^{(m)}\Big) &= 2(\sigma_g^{(m)})^4 A_q^{(m)} \\ &\quad + 4(\sigma_g^{(m)})^2 \boldsymbol{\mu}_g^{(m)\top}\mathbf{K}_q^{(m)}\mathbf{P}^{(m)}\mathbf{K}_q^{(m)}\boldsymbol{\mu}_g^{(m)}.\end{aligned} \tag{S20}$$

DeSPA treats the null-model residual-variance estimate as a plug-in quantity. It further replaces the unobserved spatial mean quadratic form in Eq. (S20) by the observable residual quadratic form

$$\mathbf{r}_g^{(m)\top}\mathbf{K}_q^{(m)}\mathbf{P}^{(m)}\mathbf{K}_q^{(m)}\mathbf{r}_g^{(m)}.$$

Because the denominator $A_q^{(m)}$ is fixed for a given slice and kernel, the resulting variance approximation for Eq. (S11) is

$$\widehat{v}_{g,q}^{(m)} = \frac{4\left(\widehat{\sigma}_g^{(m)}\right)^2\left(\mathbf{r}_g^{(m)\top}\mathbf{K}_q^{(m)}\mathbf{P}^{(m)}\mathbf{K}_q^{(m)}\mathbf{r}_g^{(m)}\right) + 2\left(\widehat{\sigma}_g^{(m)}\right)^4 A_q^{(m)}}{\left(A_q^{(m)}\right)^2}. \tag{S21}$$

The first term captures the interaction between residual noise and the realized spatial component, whereas the second term is the quadratic-form variance contributed by residual noise alone.

For two independently analyzed slices $m$ and $n$, the kernel-specific null hypothesis is $H_0 : \tau_{g,q}^{(m)} = \tau_{g,q}^{(n)}$. Under this null, the variance of the difference between the two slice-specific estimators is approximated by the sum of their variances. The standardized statistic is therefore

$$Z_{g,q}^{(m,n)} = \frac{\widehat{\tau}_{g,q}^{(m)} - \widehat{\tau}_{g,q}^{(n)}}{\sqrt{\widehat{v}_{g,q}^{(m)} + \widehat{v}_{g,q}^{(n)}}}, \tag{S22}$$

which is evaluated using the score-based standard normal approximation. The corresponding two-sided kernel-specific $p$-value is

$$p_{g,q}^{(m,n)} = 2\Phi\Big(-\left|Z_{g,q}^{(m,n)}\right|\Big), \tag{S23}$$

where $\Phi(\cdot)$ is the cumulative distribution function of the standard normal distribution. The subsequent ACAT, Fisher, and Benjamini–Hochberg combination steps are applied as described in the main Methods.

### Supplementary Tables

**Supplementary Table 1: Hardware configuration for computational analyses**

| Component | Specification |
|---|---|
| CPU | Intel Xeon Gold 6226R (2.90 GHz, 16 cores per job) |
| GPU | NVIDIA A30 (24GB GDDR6, CUDA 12.2) |
| Memory | 1TB total RAM with dynamic allocation |
| Storage | High-speed SSD |
| OS & Scheduler | Ubuntu 22.04.4 LTS with SLURM |

**Supplementary Table 2: Cell-type and cell-state abbreviations used in Figure 5.**

| Abbreviation | Full name or description |
|---|---|
| CMPN | Cholinergic, monoaminergic and peptidergic neuron |
| CP | Choroid plexus |
| dpEX | Dorsal pallium excitatory neuron |
| mpEX | Medial pallium excitatory neuron |
| mpIN | Medial pallium inhibitory neuron |
| nptxEX | Nptx$^{+}$ lateral pallium excitatory neuron |
| cckIN | Cck$^{+}$ inhibitory neuron |
| npyIN | Npy$^{+}$ inhibitory neuron |
| ntng1IN | Ntng1$^{+}$ inhibitory neuron |
| scgnIN | Scgn$^{+}$ inhibitory neuron |
| sstIN | Sst$^{+}$ inhibitory neuron |
| MSN | Medium spiny neuron |
| DMIN | Dopaminergic periglomerular inhibitory neuron |
| MCG | Microglial cell |
| Oligo | Oligodendrocyte |
| VLMC | Vascular leptomeningeal cell |
| dEGC | Developmental ependymoglial cell |
| reaEGC | Reactive ependymoglial cell |
| ribEGC | Ribosomal ependymoglial cell |
| sfrpEGC | Sfrp$^{+}$ ependymoglial cell |
| wntEGC | Wnt$^{+}$ ependymoglial cell |
| dNBL1–5 | Developmental neuroblast subclusters 1–5 |
| tlNBL | Telencephalon neuroblast |
| obNBL | Olfactory bulb neuroblast |
| rIPC1, rIPC2, rIPC4 | Regeneration intermediate progenitor cell subclusters 1, 2 and 4, respectively |
| IMN | Immature neuron |
| WSN | Wound-stimulated neuron |
| Immature (prefix) | Immature population of the corresponding neuronal type |
| Unknown | Cells of undetermined identity |

### Supplementary Figures

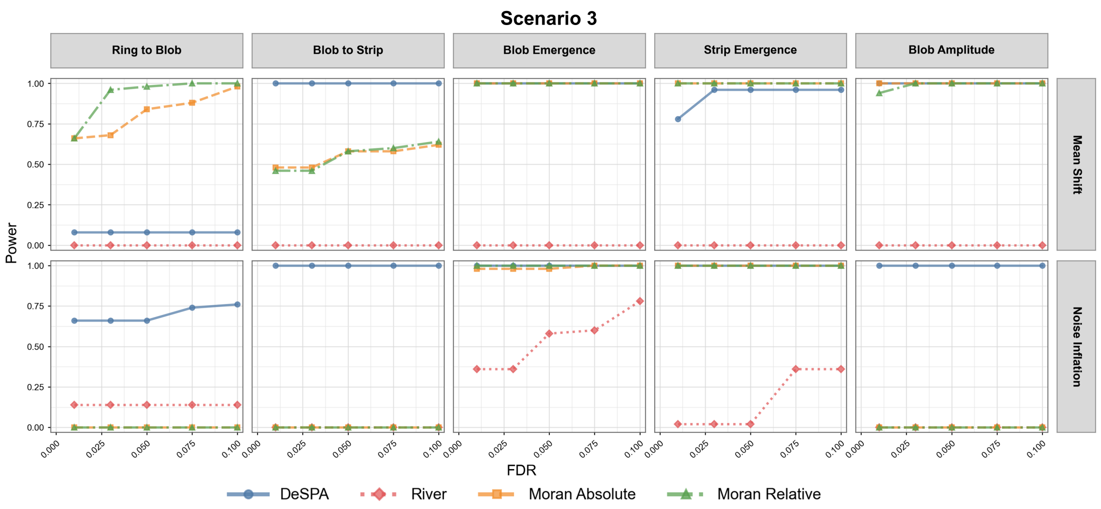


**Supplementary Figure 1: Power curves for Scenario 3. Columns correspond to the five positive spatial-pattern settings, and rows correspond to the two interference settings, mean shift and noise inflation.**

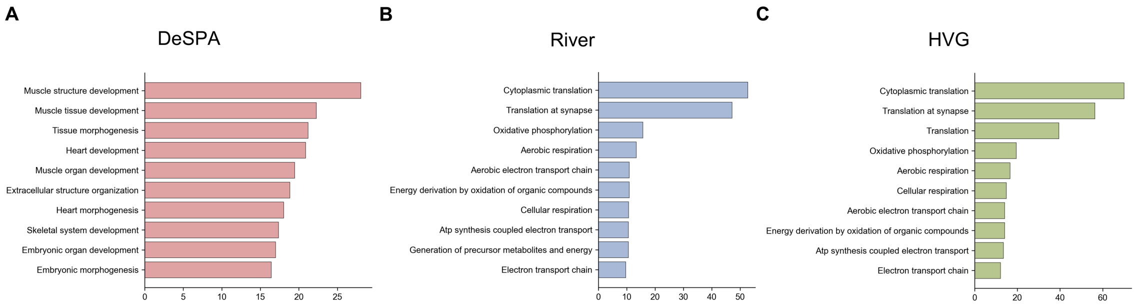


**Supplementary Figure 2: (A) GO Biological Process enrichment of the top 500 genes selected by DeSPA. (B) GO Biological Process enrichment of the top 500 genes selected by River. (C) GO Biological Process enrichment of the top 500 genes selected by HVG.**

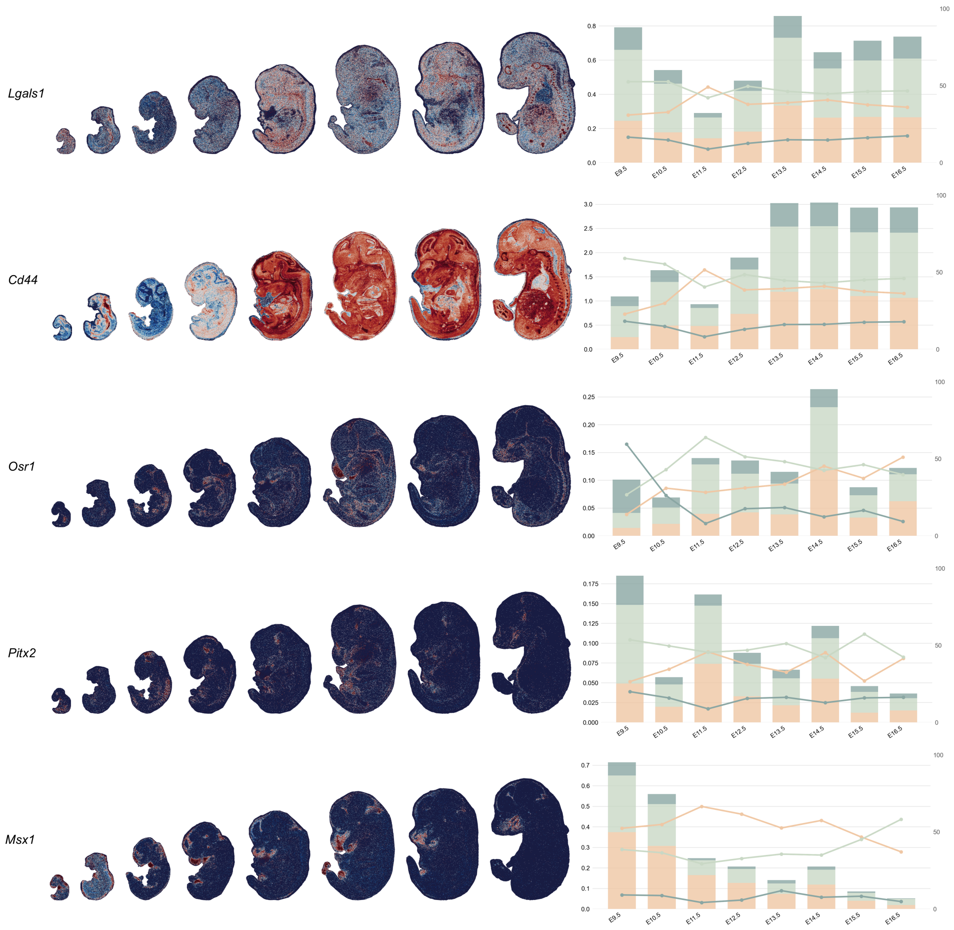


**Supplementary Figure 3: Spatial expression patterns and boundary-to-core profiles of representative developmental genes. For each gene (*Lgals1*, *Cd44*, *Osr1*, *Pitx2*, and *Msx1*), spatial expression maps from E9.5 to E16.5 are shown on the left. On the right, stacked bars show the stage-specific expression contributions from the boundary, transition, and core regions (left axis), and the overlaid lines show the same regional contributions as percentages of total gene expression (right axis).**

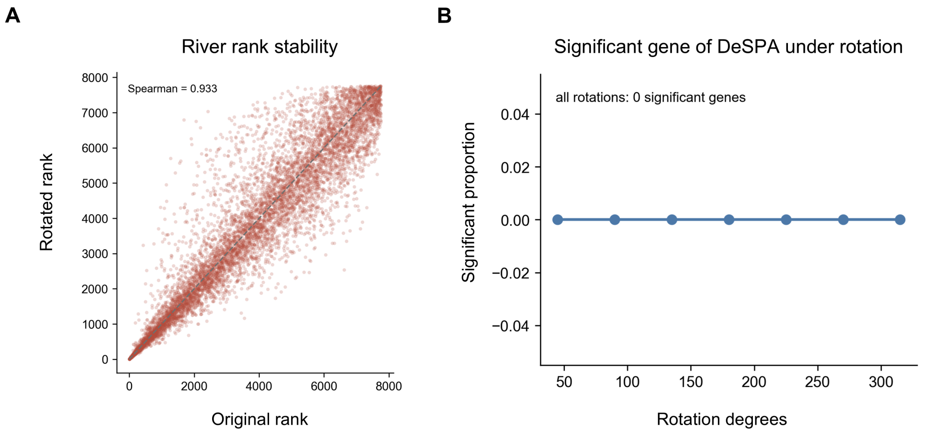


**Supplementary Figure 4: (A) Rotation analysis showing top-500 gene overlap and rank stability after rotating diabetic slices. (B) Proportion of significant genes detected by DeSPA in same-condition negative-control comparisons after rotating the spatial coordinates by $45^\circ$ increments from $45^\circ$ to $315^\circ$. No significant genes were detected at any rotation angle.**

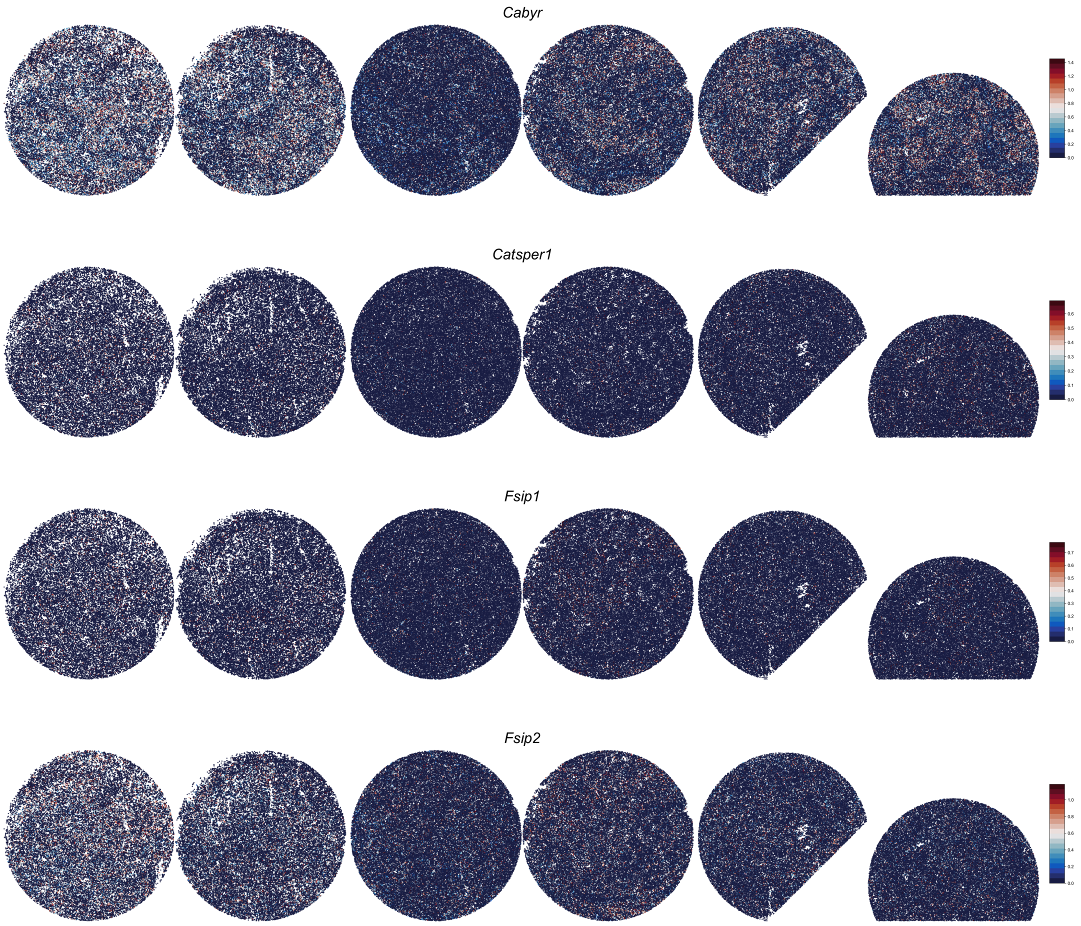


**Supplementary Figure 5: Spatial expression patterns of representative genes, including *Cabyr*, *Catsper1*, *Fsip1*, and *Fsip2* across diabetic and WT sections.**

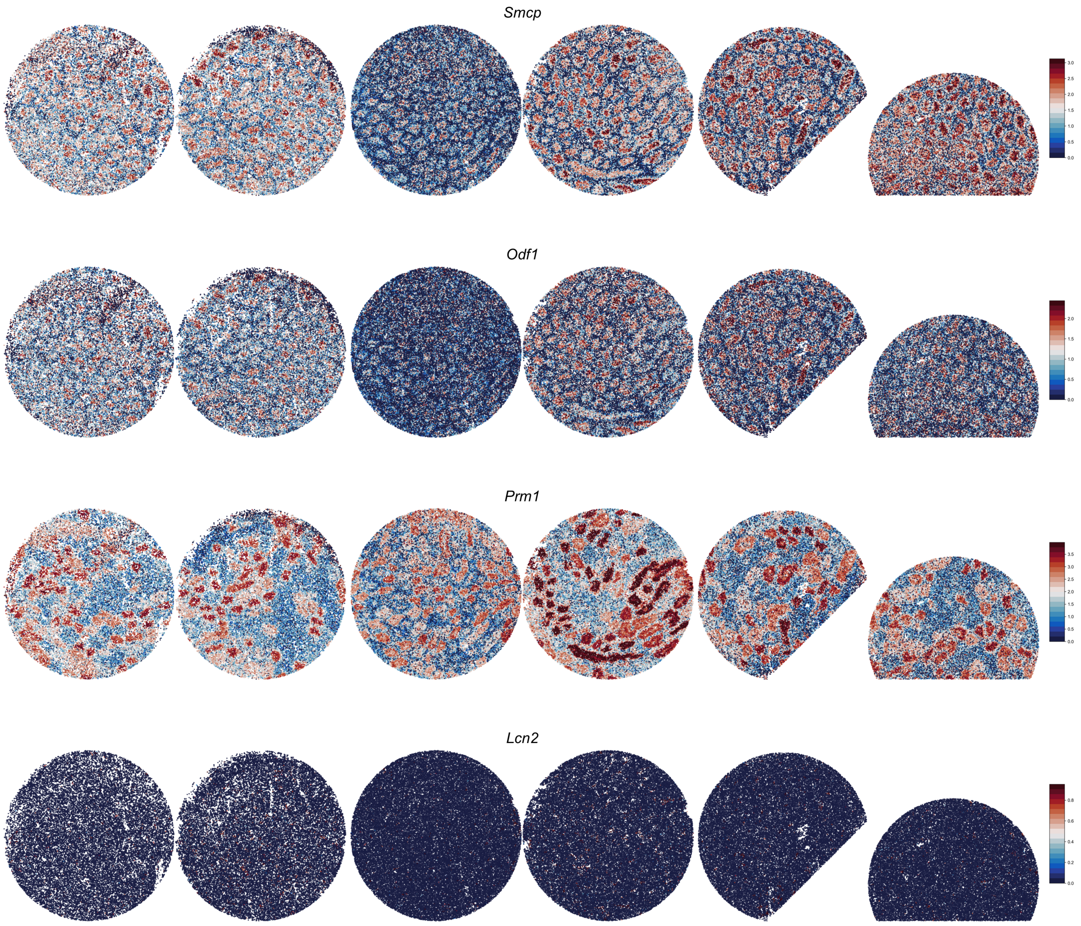


**Supplementary Figure 6: Spatial expression patterns of representative genes, including *Smcp*, *Odf1*, *Prm1*, and *Lcn2* across diabetic and WT sections.**

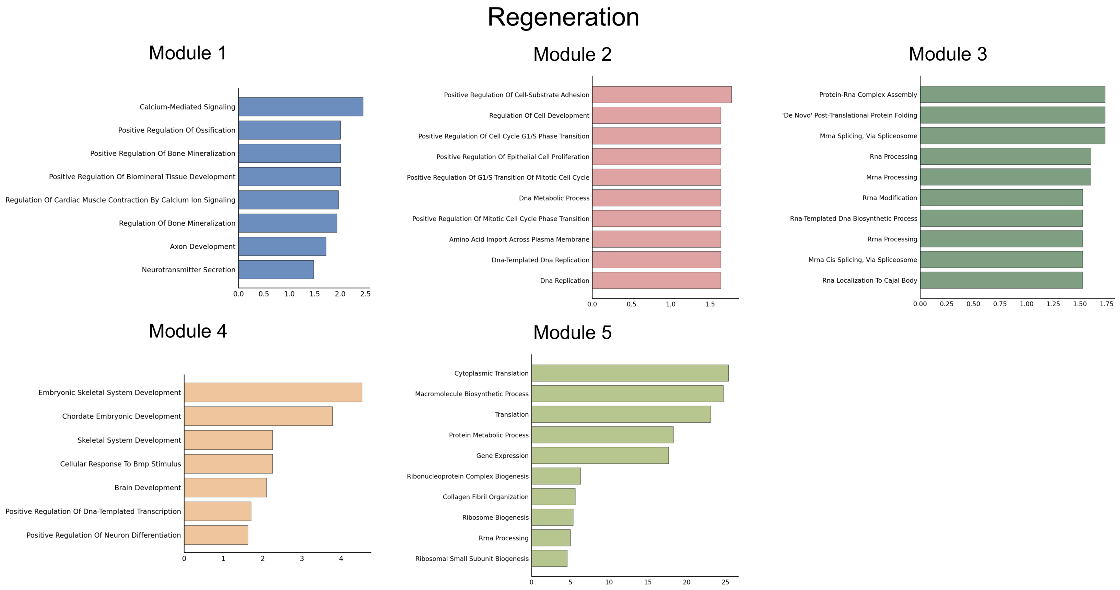


**Supplementary Figure 7: GO Biological Process enrichment of axolotl brain regeneration spatial co-expression modules.**

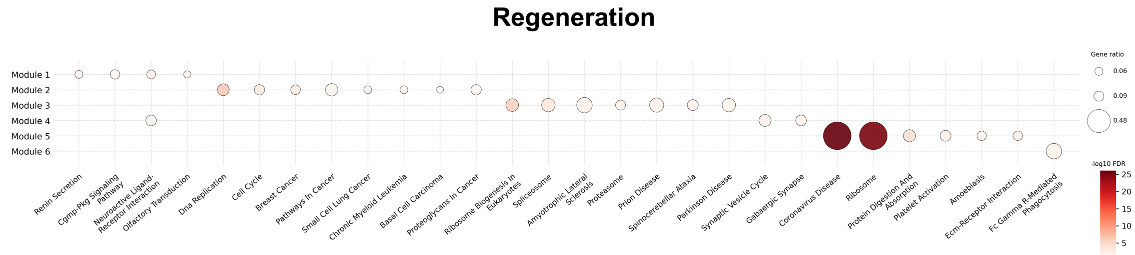


**Supplementary Figure 8: KEGG enrichment of axolotl brain regeneration spatial co-expression modules.**

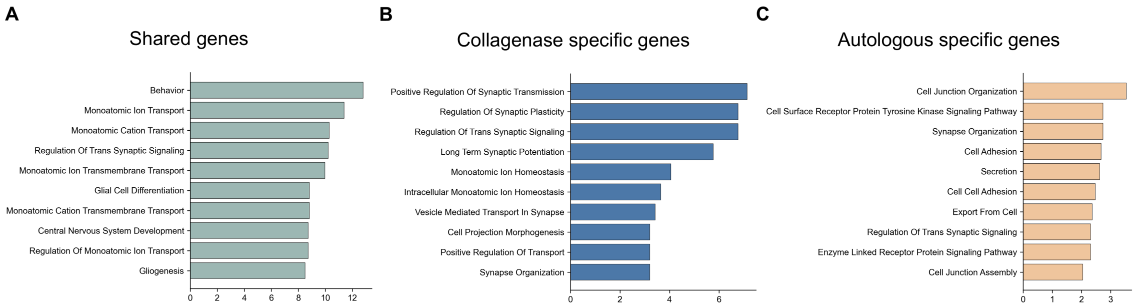


**Supplementary Figure 9: (A) GO Biological Process enrichment of shared genes. (B) GO Biological Process enrichment of collagenase specific genes. (C) GO Biological Process enrichment of autologous specific genes.**

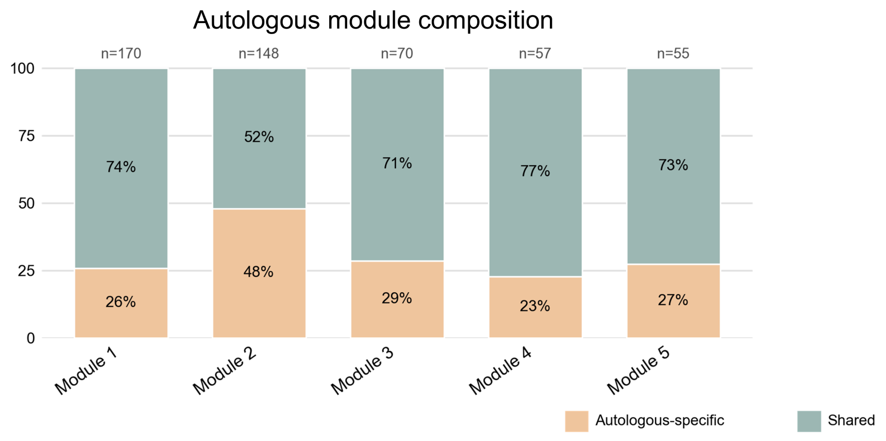


**Supplementary Figure 10: Composition of autologous-blood modules according to shared and model-specific gene membership.**

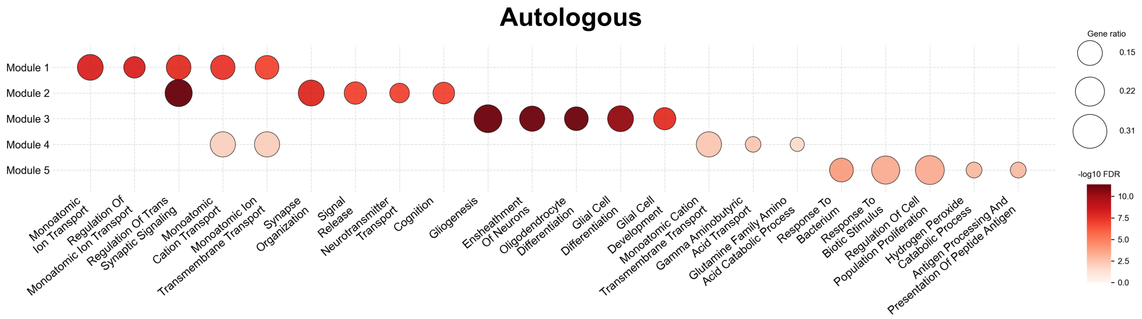


**Supplementary Figure 11: GO Biological Process enrichment of autologous spatial co-expression modules.**
